# MicroRNA-34a suppresses KLF2 to promote pathological angiogenesis through the CXCR4/CXCL12 pathway in age-related macular degeneration

**DOI:** 10.1101/2025.02.12.637499

**Authors:** Jason J. Colasanti, Joseph B. Lin, Ryo Terao, Tae Jun Lee, Andrea Santeford, Rajendra S. Apte

**Affiliations:** John F. Hardesty, MD Department of Ophthalmology & Visual Sciences, Washington University School of Medicine; St. Louis MO 63110, USA; Molecular Cell Biology Graduate Program, Roy and Diana Vagelos Division of Biology and Biomedical Sciences, Washington University School of Medicine; St. Louis MO 63110, USA; Neurosciences Graduate Program, Roy and Diana Vagelos Division of Biology and Biomedical Sciences, Washington University School of Medicine; St. Louis MO 63110, USA; Department of Ophthalmology, Graduate School of Medicine, the University of Tokyo; Tokyo 113-8655, Japan; Developmental, Regenerative, & Stem Cell Biology Graduate Program, Roy and Diana Vagelos Division of Biological and Biomedical Sciences, Washington University School of Medicine; St. Louis MO 63110, USA; Department of Developmental Biology, Washington University School of Medicine; St. Louis MO 63110, USA; Center of Regenerative Medicine, Washington University School of Medicine; St. Louis MO 63110, USA; Department of Medicine, Washington University School of Medicine; St. Louis MO 63110, USA

## Abstract

Age-related macular degeneration (AMD), characterized by pathologic choroidal neovascularization (CNV), is a leading cause of vision loss in the elderly. Vascular endothelial growth factor A (VEGFa) antagonists can prevent acute vision loss, but high treatment burden and loss of efficacy with chronic therapy highlight the need to explore alternative mechanisms. Recently, microRNA-34a (miR-34a) has emerged as a key regulator in aging and age-related diseases, but its role in neovascular AMD is unclear. In an injury-induced murine CNV model, we discovered miR-34a promoted pathological angiogenesis, without altering expression of Vegfa or its receptor Kdr, the canonical regulators of CNV. Mechanistically, miR-34a directly targets and inhibits the transcription factor KLF2 thereby upregulating the pro-angiogenic factors CXCR4 and CXCL12. Finally, we show miR-34a exacerbates CNV in aged mice and is expressed in CNV lesions excised from wet AMD patients. These findings establish a causal link between the age-related miR-34a and neovascularization in AMD.

**Teaser:** Identification of a molecular mechanism involved in the pathogenesis of a prevalent and debilitating age-related ocular disease.

## Introduction

Age-related macular degeneration (AMD), the leading cause of blindness in people over 60 years of age in the industrialized world, affects over 10 million people in the United States and over 190 million globally (*1*, *2*). There are several risk factors for AMD including smoking, high blood pressure, and obesity; however, the greatest risk factor is advanced age (*3*). In the initial stages of disease, AMD is characterized by lipoprotein-rich extracellular deposits underneath the neurosensory retina (dry AMD). Vision loss in AMD typically occurs when disease progresses to the more advanced stages characterized by either atrophic neurodegeneration (advanced dry AMD) or pathologic neovascularization (wet AMD), also called choroidal neovascularization (CNV). AMD targets the macula, the critical cone-rich region essential for the sharp central vision needed for reading, driving, and recognizing faces. The atrophic (dry), advanced dry, or advanced exudative (wet) AMD represent a disease continuum and can exist simultaneously in the same eye. Although wet AMD is less common and comprises 10-15% of disease burden, it is the most severe form and accounts for approximately 90% of severe vision loss or blindness in AMD patients (*4*). A hallmark of wet AMD is the aberrant growth of pathogenic blood vessels (i.e., CNV) underneath the retina, that ultimately results in vision loss from hemorrhage, photoreceptor loss, and fibrosis.

Vascular endothelial growth factor A (VEGFa) is a critical mediator in the development and proliferation of CNV. Based on efficacy data from randomized clinical trials, anti-VEGFa therapy is the first-line approach in the clinical management of patients affected by wet AMD (*5*). This therapeutic approach effectively blocks the interaction between VEGFa and its receptor VEGFR2 (KDR), which promotes proliferative angiogenesis. Although these therapies prevent severe vision loss in the short term and offer the possibility of improved visual acuity, treatment burden is high as multiple intraocular injections are required. Moreover, therapeutic efficacy diminishes with chronic treatment despite multiple intraocular administrations (*6–11*). Possible reasons for the decline in long-term efficacy of anti-VEGFa therapy is the existence of additional pathways influencing aberrant vascular proliferation that become dysregulated with advanced age and progressive neurodegeneration. Thus, there is an unmet need to investigate additional mechanisms of pathological angiogenesis independent of VEGF-regulated pathways, that drive age-related eye diseases such as AMD.

MicroRNAs (miRNAs) are a class of small, non-coding RNA that regulate gene expression through mRNA degradation and translational repression. Recently, miRNAs have been demonstrated to influence the molecular pathogenesis of aging (*12*) and diverse diseases including cardiovascular disease (*13*), neurodegeneration (*14*), and cancer (*15*). Indeed, our lab has previously shown that microRNAs can regulate pathological angiogenesis as seen in AMD by regulating cholesterol and fatty acid metabolism in myeloid cells (*16*, *17*). One miRNA in particular, miR-34a, contains a highly conserved seed sequence, the expression of which increases with age in rodents and nematodes, while loss of function increases median lifespan by nearly 50% in the latter (*18*).

Additionally, miR-34a is known to exacerbate disease pathology in several mouse models of age-related or age-associated diseases ranging from cardiovascular disease (*19*), Alzheimer’s disease (*20*), obesity (*21*, *22*), atherosclerosis (*23*), non-alcoholic fatty liver disease (*24*), lung fibrosis (*25*), and diabetes (*26*). Although miR-34a is upregulated in the ocular tissues of AMD patients (*27*, *28*) little is known regarding its molecular role in AMD progression. Given its multifaceted role in vascular endothelial cell dysfunction (*29*, *30*), miR-34a is a prime candidate for investigating molecular mechanisms of age-related pathological angiogenesis in the eye.

Here, we investigated miR-34a in the context of neovascular AMD using a well-established mouse model of laser injury induced CNV (*31*). We characterized for the first time how miR-34a expression is dysregulated in murine choroidal vasculature following laser injury and demonstrated the cell type where its expression directly influences CNV. By combining bulk and single-cell RNA sequencing data, we identified a previously unknown target of miR-34a in vascular endothelial cells. Using both *in vitro* and *in vivo* methods, we validated this target of miR-34a was dysregulated in vascular endothelial cells without altering VEGFa or KDR expression. We investigated miR-34a in the context of murine ocular aging and found that it also exacerbates CNV in aged mice. Finally, we showed that miR-34a is upregulated in human CNV, and its downstream factors are dysregulated in human vascular endothelial cells in the choroid with age or in neovascular AMD. This study unveiled a molecular pathway through which miR-34a governs pathological angiogenesis among individuals with wet AMD in a VEGFa independent manner. These results could contribute to the creation of alternative therapeutic approaches, augmenting the current anti-VEGFa treatments by focusing on additional factors that facilitate anomalous vascular proliferation.

## Results

### MiR-34a is upregulated in choroidal vascular endothelial cells after injury

To evaluate ocular miR-34a expression during pathological angiogenesis, we induced CNV lesions in mice via a laser injury model (Fig. 1A). We isolated microRNA from retinal pigment epithelium (RPE)/choroidal cell lysates at seven days post-injury, corresponding to the point at which CNV lesions attained their maximal sizes. We then performed quantitative PCR (qPCR) and found a statistically significant increase of miR-34a transcript levels in RPE/choroid lysates seven days after laser injury compared to lysates from uninjured samples (Fig. 1B), corroborating previous findings (*32*). To evaluate the temporal course of miR-34a activation in murine CNV, we visualized miR-34a expression through in-situ hybridization staining of murine whole-eye sections that were collected over several days following laser injury. We found there was a statistically significant increase in miR-34a expression within CNV lesions over time (Fig. 1, C and D). Since laser CNV lesions consist of endothelial cell populations and immune cell populations (*17*, *33*, *34*) (Fig. S1A), we sought to determine which cell type most closely co-localized with miR-34a expression. We found that miR-34a expression strongly overlapped with CD31 protein, a marker for vascular endothelial cells (Fig. 1E). In contrast, we did not observe a strong overlap between miR-34a expression and known immune cell markers (Fig. S1, B to D). In addition to choroidal vascular endothelial cells, we also found that miR-34a was consistently expressed in a subset of retinal ganglion cells (RGCs) regardless of injury status (Fig. S2), possibly due to its roles in neuronal development and function (*35*, *36*).

**Fig. 1.**
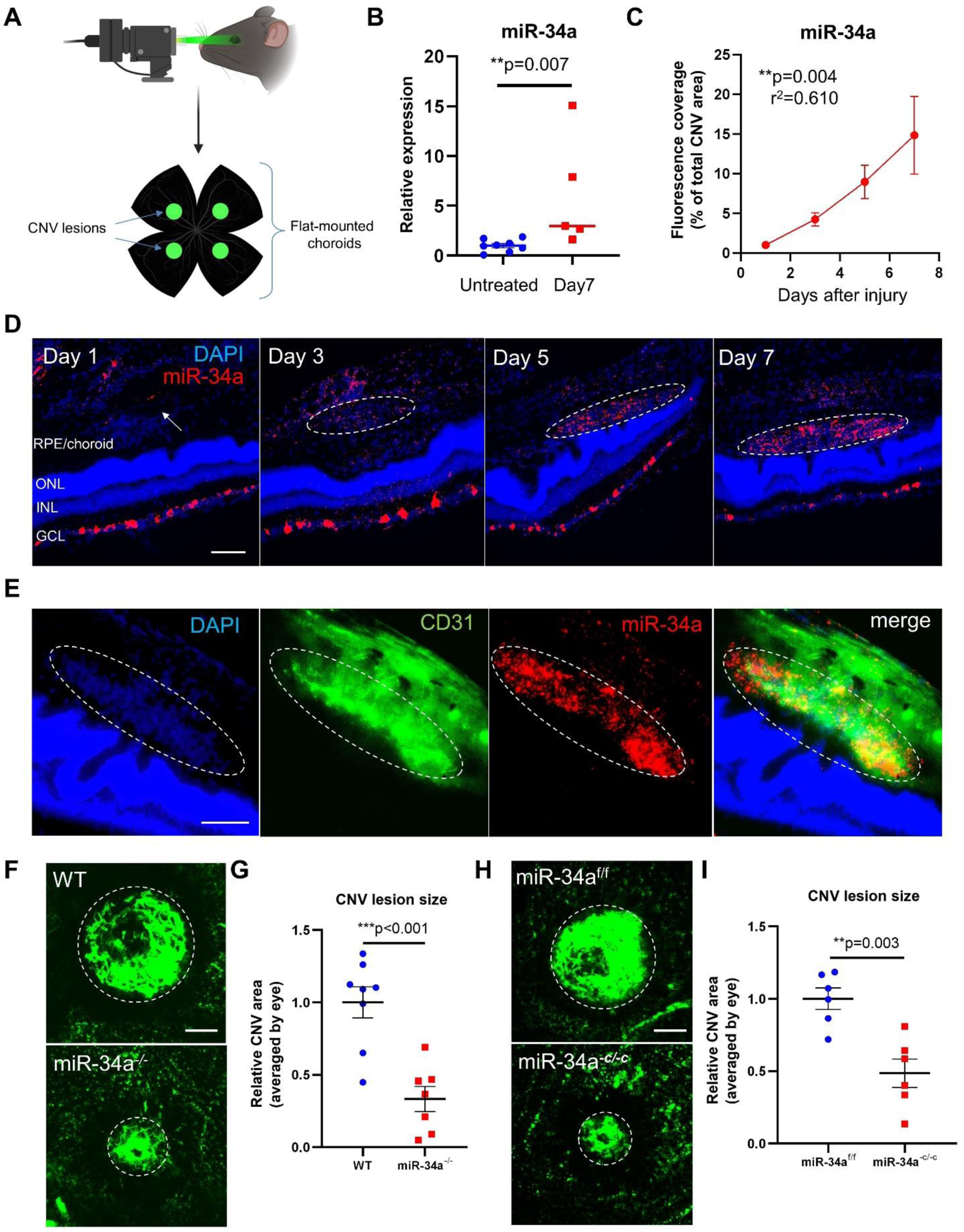
MiR-34a is upregulated in choroidal vascular endothelial cells after laser injury and promotes pathological angiogenesis. **(A)** A schematic depicting the laser-induced CNV mouse model. Created with BioRender.com (agreement number: IU277NSX86). **(B)** Relative expression of miR-34a was increased in the RPE/choroid following laser injury as measured by qPCR (U6 used as an endogenous control, n = 5-8 mice/group, Mann-Whitney test). **(C)** Quantification of the relative fluorescence intensity of miR-34a staining in CNV lesions collected on multiple days following laser injury (n = 3-4 mice/group, One-way ANOVA, test for linear trend). **(D)** Images of whole-eye sections from WT mice on days 1, 3, 5, and 7 post-laser injury. ONL denotes outer nuclear layer, INL denotes inner nuclear layer, and GCL denotes ganglion cell layer. **(E)** Images of whole-eye sections dual-stained via immunohistochemistry for CD31 and via in situ hybridization for miR-34a. **(F)** Flat mount images of CNV lesion sizes between WT and whole-body miR-34a^-/-^ mice. **(G)** Quantification of CNV lesions sizes between WT and miR-34a^-/-^ mice (n = 7-8 mice/group, unpaired Welch’s t test). **(H)** Images of CNV lesion sizes between miR-34a^fl/fl^ (miR-34a^f/f^) and miR-34a^-cdh5/-cdh5^ (miR-34a^-c/-c^) mice. **(I)** Quantification of CNV lesion sizes between miR-34a^fl/fl^ and miR-34a^-cdh5/-cdh5^ mice (n = 4 mice/group, unpaired Welch’s t test). Scale bars = 100µm.

### MiR-34a promotes pathological angiogenesis

To determine whether there was a causal relationship between miR-34a and pathological angiogenesis, we induced CNV in mice that were globally deficient for miR-34a (miR-34a^-/-^). At baseline, we did not observe any ocular abnormalities in miR-34a^-/-^ mice via fundoscopic examination (Fig. S3A), consistent with previous reports indicating normal gross anatomical features of miR-34a^-/-^ mice (*37*, *38*). Additionally, miR-34a^-/-^ mice had normal photoreceptor function compared to wild-type (WT) mice as assessed by electroretinograms (ERG) (Fig. S3B). By contrast, after laser-induced injury CNV lesions in miR-34a^-/-^ mice were significantly reduced compared to WT controls (Fig. 1, F and G). Because our immunohistochemistry staining suggested that miR-34a was expressed primarily by vascular endothelial cells in the outer retina and RGCs in the inner retina (Fig. 1E and fig. S2A), we sought to determine the role of miR-34a action in these cell types. To do this, we generated mice that were deficient in miR-34a in either vascular endothelial cells or in RGCs. We crossed mice with miR-34a flanked by loxP sites (*35*) (miR-34a*^fl/fl^*) with either mice harboring vascular endothelial-specific Cre or mice harboring RGC-specific Cre to generate mice lacking miR-34a specifically in either vascular endothelial cells (*39*) (miR-34a^-*cdh5/cdh5*^) or RGCs (*40*) (miR-34a^-*vglut2/-vglut2*^) and validated that miR-34a was successfully knocked-out in the respective cell types (Fig. S4A). Then, we repeated the CNV lesion size experiment and observed significantly reduced CNV lesions in miR-34a*^-cdh5/-cdh5^* mice compared to controls (Fig. 1, H and I), suggesting that miR-34a promotes pathological angiogenesis via vascular endothelial cells. However, we did not observe a significant difference in CNV lesions in miR-34a^-^ *^vglut2/-vglut2^* mice compared to controls (Fig. S5, A and B), consistent with our finding that RGC expression of miR-34a is independent of CNV status and further suggesting that miR-34a expression in RGCs may not play a meaningful role in pathological angiogenesis. Because RGC expression of miR-34a was independent of CNV status, we did not study RGC-expressed miR-34a further in this work, though further studies on miR-34a in these cells may be warranted. Taken together, our data indicate that miR-34a is activated in vascular endothelial cells to promote pathological angiogenesis in the eye.

### MiR-34a promotes pathological angiogenesis without altering expression of VEGFa or KDR

Given that the VEGFa/KDR axis represents the most extensively studied mechanism for disease-associated vascular proliferation in the eye and the major canonical driver of CNV, we next assessed the impact of miR-34a on the expression of either *VEGFa* or *KDR*. To model the vascular endothelial cells in CNV, we used human umbilical vein endothelial cells (HUVECs), as utilized in several previous studies (*41–47*). First, we confirmed the successful transfection of miR-34a mimics into HUVECs by comparing the relative miR-34a expression to HUVECs transfected with scrambled control mimics (Fig. S6). Next, we transfected HUVECs with either miR-34a mimics or scrambled control mimics and measured relative transcript expression of *VEGFa* and *KDR* via qPCR (Fig. 2A). Interestingly, we did not find significant differences for either *VEGFa* or *KDR* expression between HUVECs transfected with a miR-34a mimic or treated with a scrambled control (Fig. 2, B and C). *In vivo*, we measured Vegfa and Kdr protein levels in CNV lesions and compared miR-34a^-/-^ mice to WT controls. In agreement with our HUVEC data, we also observed no differences in Vegfa or Kdr expression in CNV lesions between these two groups (Fig. 2, D to G). These findings were particularly noteworthy because they suggest that miR-34a promotes pathological angiogenesis in the eye through a mechanism other than the VEGFa/KDR axis.

**Fig. 2.**
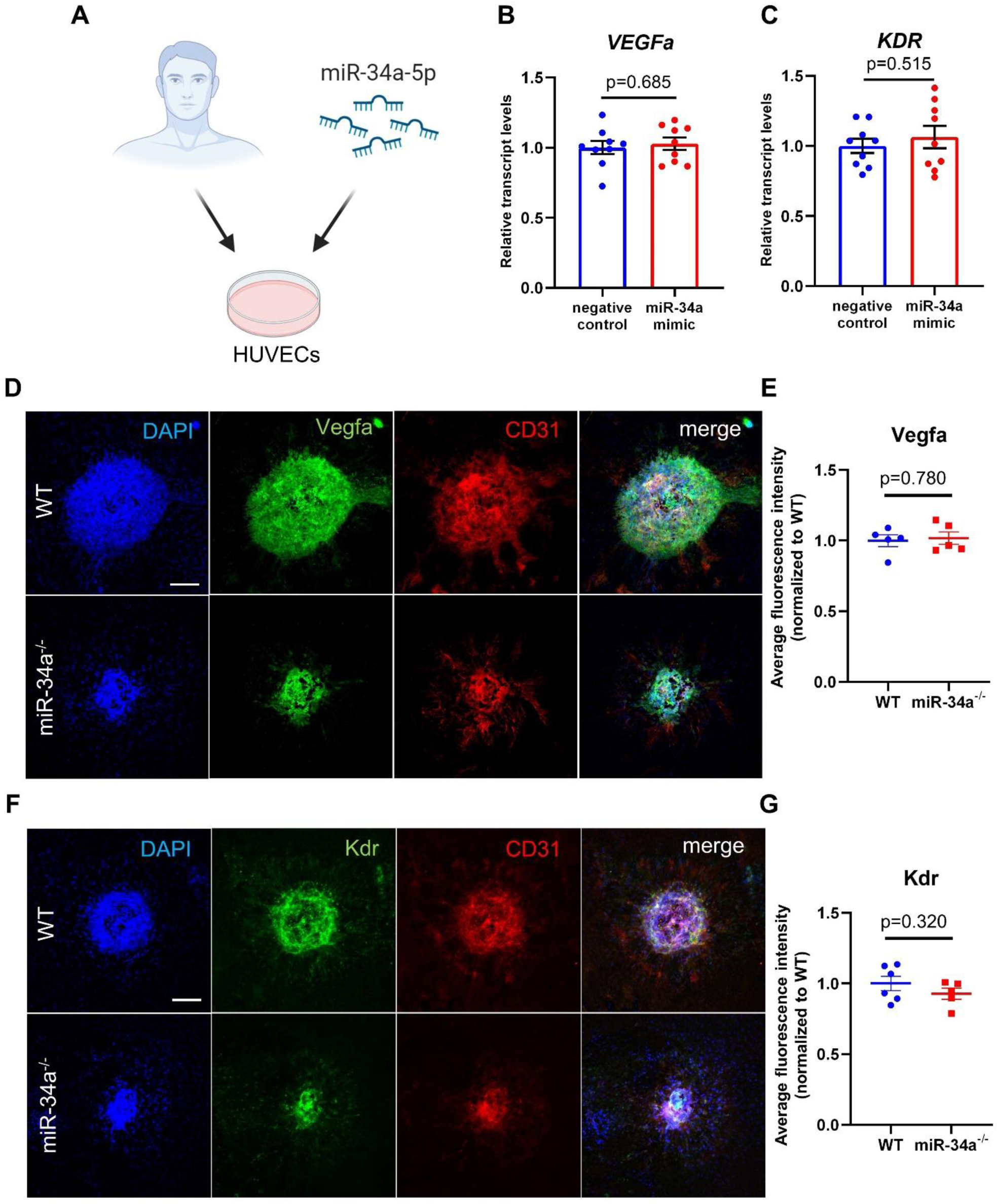
MiR-34a does not alter VEGFa or KDR expression in vascular endothelial cells. **(A)** Schematic depicting human-derived cells (HUVECs) being treated with miR-34a-5p mimics in cell culture. Created with BioRender.com (agreement number: UO277NTFVP). **(B, C)** HUVECs transfected with a miR-34a mimic did not have significant differences in *VEGFa* or *KDR* relative transcript expression via qPCR (actin used as an endogenous control, n = 9 replicates/group, unpaired Welch’s t test). **(D, F)** Images of murine CNV lesions stained with Vegfa or Kdr and vascular endothelial cell marker CD31. **(E, G)** Quantification of relative fluorescence intensity for vegfa and kdr stained murine CNV lesions (n = 5-6 mice/group, unpaired Welch’s t test). Scale bars = 100µm.

### KLF2 is a direct target of miR-34a in vascular endothelial cells

To uncover a molecular mechanism for how miR-34a promotes vascular proliferation in the eye, we used an integrative transcriptomic approach to identify targets of miR-34a in CNV. We transfected HUVECs with a miR-34a mimic and performed RNA-sequencing with differential gene expression analysis via the Limma (*48*) package for R. To determine the direct targets of miR-34a, we identified the 1,919 genes that were significantly downregulated (adj. p<0.05, |fc|>1.2) by over-expression of miR-34a. To identify which of these, if any, were relevant to CNV, we utilized a publicly available single-cell RNA sequencing dataset containing choroidal vascular endothelial cells in murine CNV lesions (*49*). We identified the overlap between the 1,919 genes downregulated by miR-34a in HUVECs with the 299 genes downregulated in murine CNV (Fig. 3A). We found 39 genes that overlapped between these two data sets, and further refined our search to seven genes by using the miRDB online database (*50*) to find predicted targets of miR-34a. Here, we found *KLF2, EDNRB, IVNS1ABP, PITPNC1, RARB, S1PR3, and SGPP1* fit all these criteria (Fig. 3A).

**Fig. 3.**
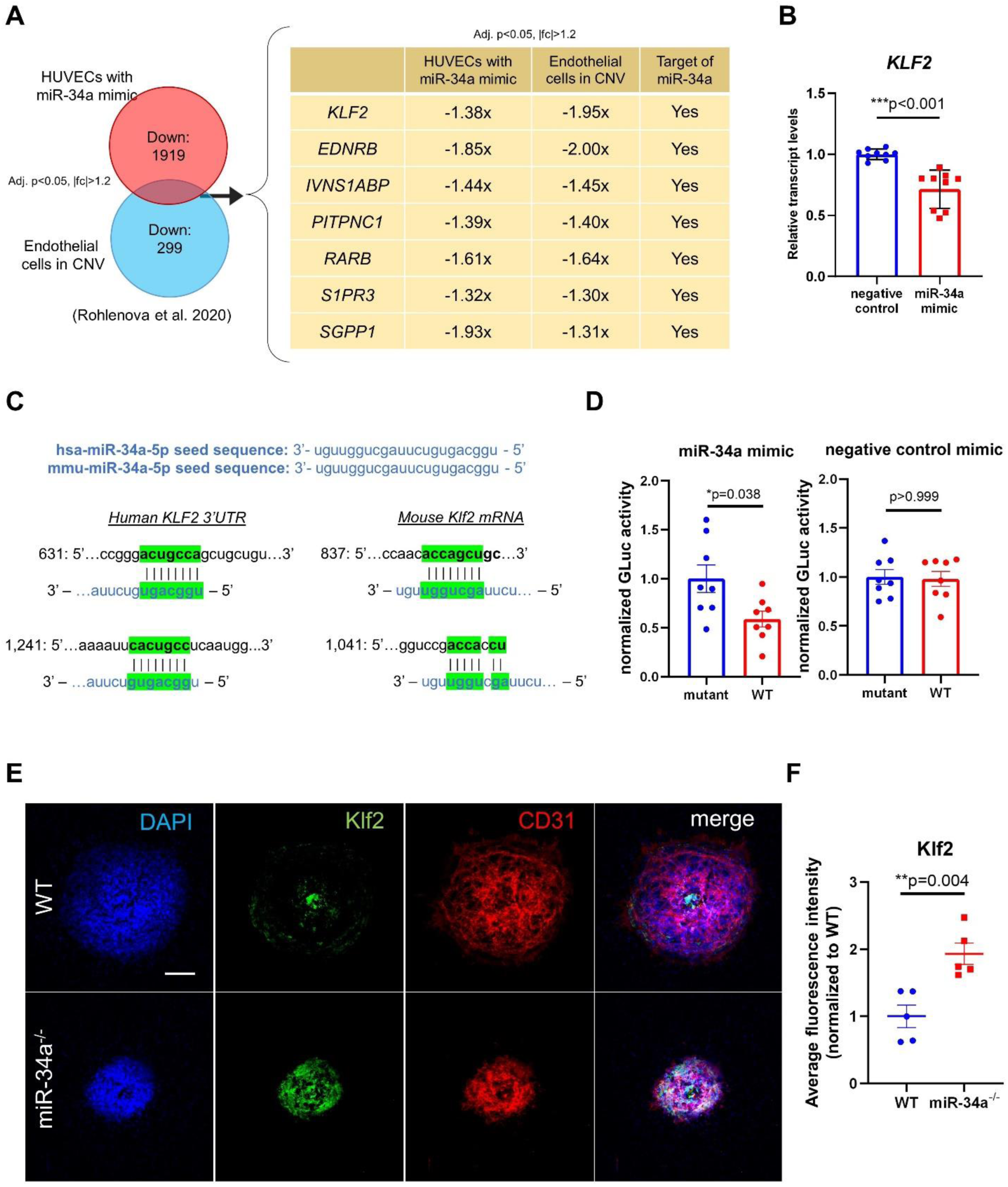
KLF2 is a direct target of miR-34a in vascular endothelial cells. **(A)** Differential gene expression analysis was performed on bulk RNA-seq data of HUVECs transfected with miR-34a mimics. Genes downregulated in HUVECs after miR-34a transfection were compared to those in a single cell RNA-seq dataset of vascular endothelial cells in murine CNV (Adj. p<0.05, |fold change| > 1.2). Seven genes that were shared between both datasets were predicted to be direct targets of miR-34a-5p. **(B)** qPCR validation of bulk-RNA seq data showing *KLF2* was downregulated after miR-34a transfection (actin used as an endogenous control, n = 9/group, unpaired Welch’s t test). **(C)** Human and mouse miR-34a-5p seed sequences are identical. Sequence overlaps between the miR-34a seed sequence with human *KLF2* 3’UTR and mouse *Klf2* mRNA. **(D)** Dual luciferase assay with a reporter plasmid containing the *KLF2* 3’UTR with the miR-34a-5p predicted binding sites either intact or removed (mutant), co-transfected with miR-34a or negative control mimics. Co-transfection with the intact *KLF2* 3’UTR and miR-34a mimic led to a decreased signal compared to the mutant plasmid (n = 8/group, unpaired Welch’s t test). **(E)** Images of murine CNV lesions stained for Klf2. **(F)** Quantification of relative fluorescence intensity for Klf2 stained murine CNV lesions (n = 5 mice/group, unpaired Welch’s t test). Scale bar = 100µm.

Of these seven predicted targets, we found Krüppel-like transcription factor 2 (*KLF2*) to be the most promising since it is widely known as a potent inhibitor of angiogenesis (*51*– *56*). Additionally, KLF2 may exert vasoprotective effects in other age-related vascular diseases including atherosclerosis (*57*) and COVID-19 (*58*). Therefore, we hypothesized that upregulation of miR-34a in vascular endothelial cells could downregulate *KLF2* and drive pathologic angiogenesis. To validate our miR-34a target screen, we confirmed that *KLF2* was indeed significantly downregulated with increased miR-34a expression in HUVECs (Fig. 3B). By comparing seed sequences of miR-34a-5p with human and mouse *KLF2* 3’UTR and mRNA, we found several sequences that either fully or partially overlapped (Fig. 3C). The direct inhibition of *KLF2* by miR-34a was validated by dual luciferase assays using the human 3’UTR of *KLF2* (Fig. 3D). *In vivo*, we observed that the expression of Klf2 was relatively low in CNV lesions of WT mice via immunohistochemistry (Fig. 3E). To determine whether miR-34a influenced Klf2 protein expression in pathological angiogenesis, we compared CNV lesions of miR-34a^-/-^ and WT mice via immunohistochemistry and found that relative Klf2 expression was significantly upregulated by almost 2-fold in the absence of miR-34a (Fig. 3, E and F). Overall, our results demonstrate KLF2 may be a direct target of miR-34a during disease-associated vascular proliferation in the eye.

### The proangiogenic factors CXCR4 and CXCL12 are upregulated by miR-34a

KLF2 is a transcription factor known to suppress the VEGFa/KDR axis in vascular endothelial cells (*51*); however, VEGFa and KDR expression were unaltered by miR-34a in our *in vitro* and *in vivo* experiments (Fig. 2, B to G). Thus, we sought an alternative mechanism by which the miR-34a/KLF2 pathway mediates angiogenesis. To accomplish this, we extensively reviewed prior literature for targets of KLF2 that also modulate CNV lesion development. Of note, KLF2 was previously reported to suppress both the C-X-C chemokine receptor type 4 (*CXCR4*) and its canonical ligand C-X-C motif chemokine 12 (*CXCL12*) in vascular endothelial cells (*59*, *60*), both of which play significant roles in blood vessel formation (*61–67*). Furthermore, both Cxcr4 and Cxcl12 are known to be upregulated in the choroid after laser-induced CNV and exacerbate pathological angiogenesis (*68–70*).

First, we assessed whether *CXCR4* and *CXCL12* were altered by miR-34a in vascular endothelial cells. Interestingly, *CXCR4* was significantly upregulated in both the miR-34a transfected HUVEC data set and the publicly available single-cell murine laser CNV dataset (Supplemental Data File 1, adj. p<0.05, |fc|>1.2). To validate our RNA-sequencing data, we treated HUVECs with a miR-34a mimic and found that *CXCR4* and *CXCL12* transcripts were both significantly increased compared to the scrambled control mimic (Fig. 4, A and B). *In vivo*, we validated that protein expression of both Cxcr4 and Cxcl12 were significantly downregulated in CNV lesions of miR-34a^-/-^ mice compared to WT (Fig. 4, C to F). These findings suggest that miR-34a indirectly upregulates the proangiogenic factors CXCR4 and CXCL12 in vascular endothelial cells.

**Fig. 4.**
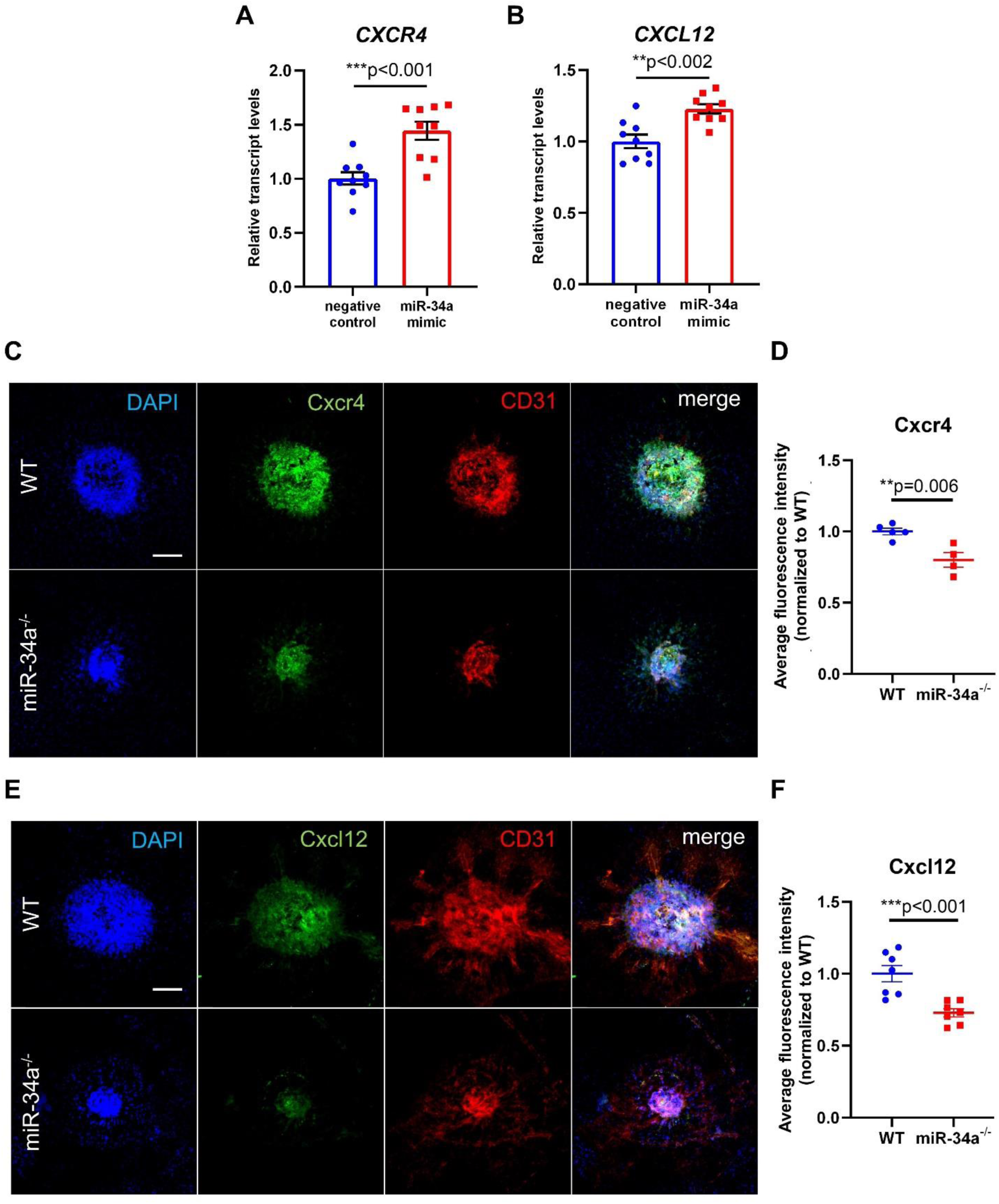
The proangiogenic factors CXCR4 and CXCL12 are upregulated by miR-34a. **(A, B)** qPCR validation of bulk-RNA seq data showing *CXCR4* and *CXCL12* are upregulated after miR-34a transfection (actin used as an endogenous control, n = 9/group, unpaired Welch’s t test). **(C, E)** Images of murine CNV lesions stained for Cxcr4 or Cxcl12. **(D, F)** Quantification of relative fluorescence intensity for Cxcr4 and Cxcl12 stained murine CNV lesions (n = 4-7 mice/group, unpaired Welch’s t test). Scale bars = 100µm.

### Suppression of KLF2 upregulates the CXCR4/CXCL12 axis and promotes pathological angiogenesis

To demonstrate the effects of *KLF2* suppression in vascular endothelial cells on gene expression, we transfected HUVECs and mouse vascular endothelial cells (MVECs) with locked nucleic acids “Gapmers” (LNA Gapmers) specifically targeting the *KLF2* sequence or scrambled sequences. After transfection, we isolated RNA and performed qPCR on several genes of interest. As expected, *KLF2* transcript expression was significantly decreased in both HUVECs and MVECs (Fig. 5, A and F). Additionally, relative transcript levels of *CXCR4* and *CXCL12* were both significantly increased when *KLF2* was suppressed (Fig. 5, B, C, G, and H). These results corroborate our earlier findings in which KLF2 expression had an inverse relationship with CXCR4 and CXCL12 expression in HUVECs transfected with miR-34a mimics and in CNV lesions between miR-34a^-/-^ and WT mice. In agreement with our earlier data, we did not observe any differences in *VEGFa* transcript expression in either HUVECs or MVECs treated with *KLF2* Gapmers compared to scrambled control Gapmers (Fig. 5, D and I). Although there was no difference for *KDR* expression in HUVECs after *KLF2* suppression (Fig. 5E), there was a significant increase in relative transcript expression for *Kdr* in MVECs (Fig. 5J). While this finding was not supported by our earlier CNV data, this may be due to an off-target sequence effect or a species-specific effect in MVECs. Next, we used these same LNA Gapmers to knock-down *Klf2* gene expression in miR-34a^-/-^ mice via daily intraperitoneal injections after laser injury, a strategy that we have previously published (*16*). CNV lesions of miR-34a^-/-^ mice treated with *Klf2* Gapmers were significantly larger than those from miR-34a^-/-^ treated with negative control Gapmers (Fig. 5, K and L). Together, these data show *KLF2* suppresses *CXCR4* and *CXCL12* without altering expression of *VEGFa* or *KDR* in human vascular endothelial cells and that KLF2 has anti-angiogenic properties *in vivo* in the laser injury-induced CNV model in miR-34a*^-/-^* mice.

**Fig. 5.**
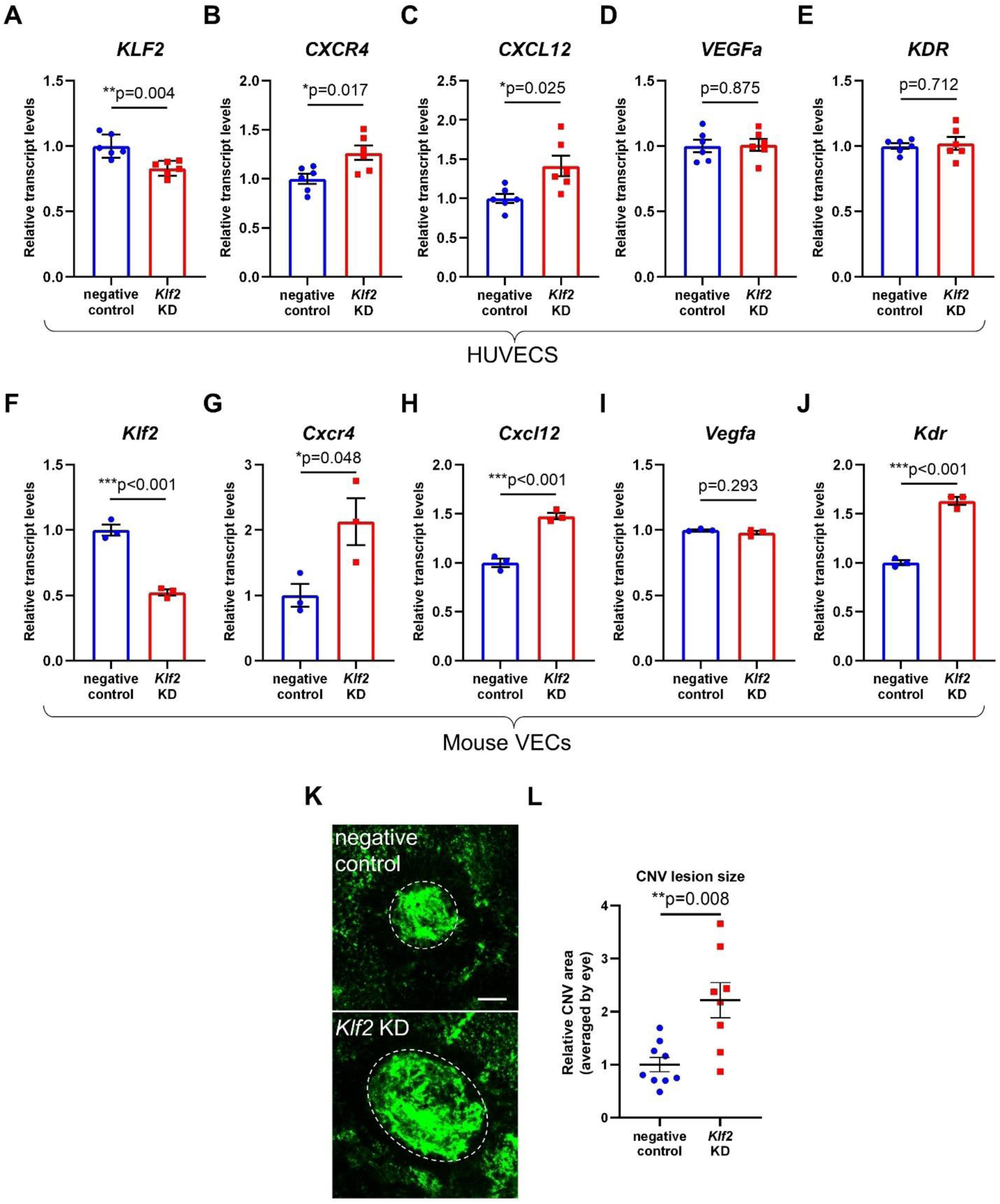
Suppression of KLF2 upregulates the CXCR4/CXCL12 axis and promotes pathological angiogenesis. **(A-E)** Relative transcript levels for *KLF2*, *CXCR4*, *CXCL12*, *VEGFa*, and *KDR* in HUVECs after transfection with either a *Klf2* Gapmer or negative control Gapmer (actin used as an endogenous control, n = 6/group, unpaired Welch’s t test). **(F-J)** Relative transcript levels for *Klf2*, *Cxcr4*, C*xcl12*, *Vegfa*, and *Kdr* in MVECs after transfection with either a *Klf2* Gapmer or negative control Gapmer (actin used as an endogenous control, n = 3/group, unpaired Welch’s t test). **(K)** Images of CNV lesion sizes between miR-34a^-/-^ mice treated with either a *Klf2* Gapmer or negative control Gapmer. **(L)** Quantification of CNV lesion sizes between miR-34a^-/-^ mice treated with either a *Klf2* Gapmer or negative control Gapmer (n = 5 mice/group, unpaired Welch’s t test) Scale bar = 100µm.

### MiR-34a promotes pathological angiogenesis in aged mice

Because miR-34a is widely implicated in aging and age-related diseases (*18–26*), we assessed whether miR-34a expression changed, or had any impact on ocular function with age. To evaluate changes in miR-34a expression with age, we sectioned whole eyes of young WT and aged WT mice and stained for miR-34a via in-situ hybridization. For this, we specifically selected sections that cut through the optic nerve so as to obtain comprehensive snapshots of the eye at the same location (Fig. S7, A and B). Once again, we only observed positive miR-34a signal in RGCs, and not in the RPE, choroid, or vasculature. To determine whether miR-34a expression was altered with age, we measured the relative fluorescence intensity of the entire ganglion cell layer in young and aged WT mice and found no significant differences (Fig. S7C). Additionally, we found the number of miR-34a-positive RGCs in young and aged mice to be relatively similar (Fig. S7D). Taken together, these data suggest that miR-34a expression by RGCs as measured via in situ hybridization does not change with age in the murine eye.

To assess the functional impacts of miR-34a during ocular aging, we aged miR-34a^-/-^ mice for 20 months and then performed several experiments while comparing to age-matched WT mice. As seen in young miR-34a^-/-^mice, we did not observe any ocular abnormalities in aged miR-34a^-/-^ mice via fundoscopy (Fig. 6A). Similarly, we did not observe any significant differences in photoreceptor function via ERG between aged miR-34a^-/-^ and WT mice (Fig. 6B), suggesting that miR-34a is not necessary for normal photoreceptor function in aged mice. To assess if miR-34a increased in CNV lesions with age, we stained ocular tissue from young and old WT mice after laser injury. We found no changes in miR-34a expression between young and old CNV (Fig. 6, C and D), possibly because miR-34a expression may become fully saturated within vascular endothelial cells seven days after laser injury. This finding may also further support the use of young mice in laser induced CNV as a robust model for phenocopying wet AMD pathology. Next, we induced CNV in aged miR-34a^-/-^ and aged WT mice. Both groups of aged mice had relatively larger CNV lesion sizes compared to young mice, as described previously (*71*). Interestingly, aged miR-34a^-/-^ mice also had significantly diminished CNV lesions sizes compared to aged WT mice (Fig. 6, E and F), suggesting that the mechanism by which miR-34a promotes pathological angiogenesis is conserved in advanced age.

**Fig. 6.**
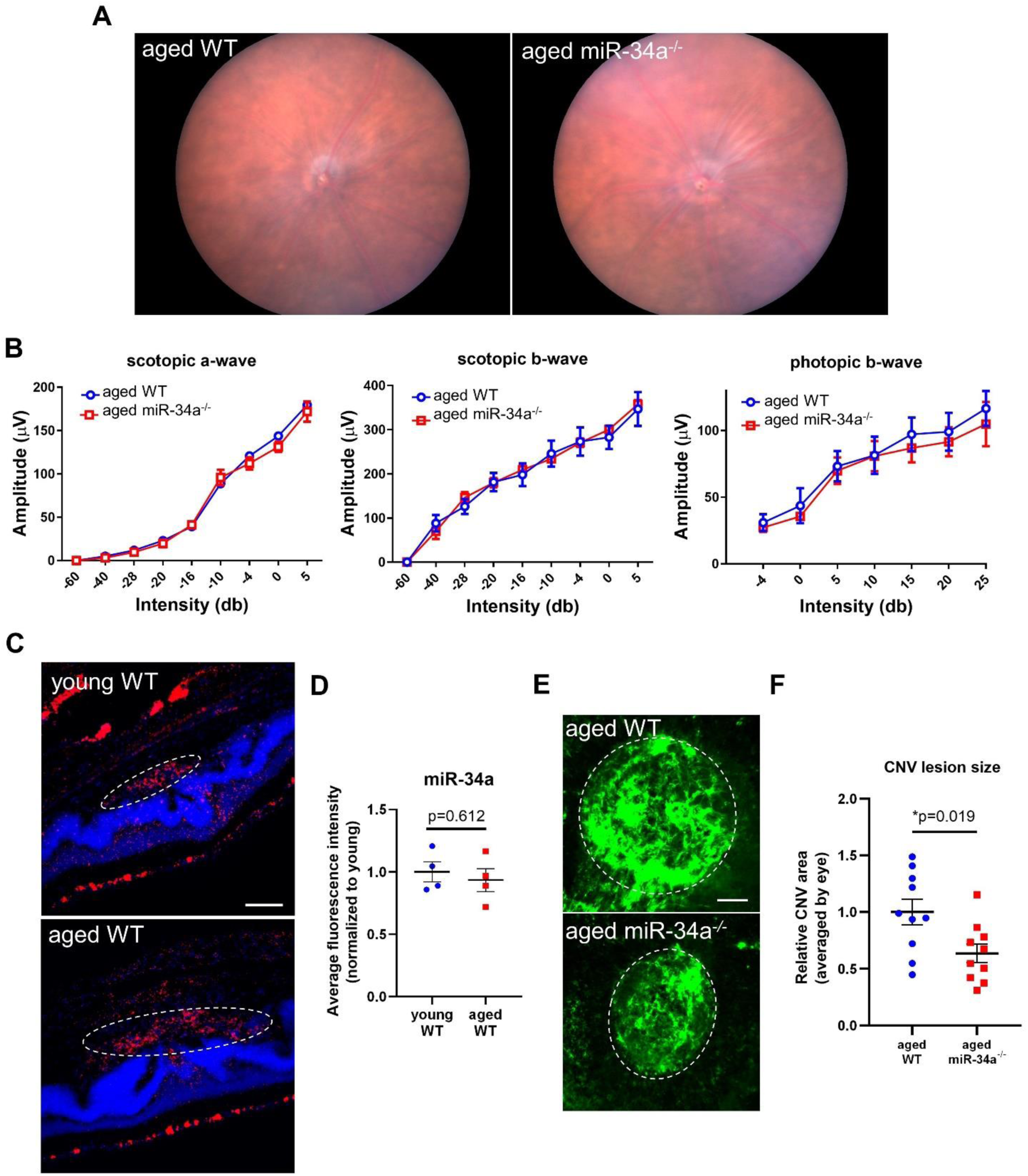
MiR-34a promotes pathological angiogenesis in aged mice. **(A)** Fundoscopy images of aged WT and miR-34a^-/-^ mice. **(B)** ERG data from aged WT and aged miR-34a^-/-^ mice (n = 4 mice/group, Two-way ANOVA with Bonferroni’s multiple comparison test). **(C)** Cross-sectional images of CNV stained via in situ hybridization for miR-34a (red) and DAPI (blue) in both young WT and aged WT mice. **(D)** Quantification of average fluorescence intensity for miR-34a staining in CNV lesions from young WT and aged WT mice (n = 4/group, unpaired Welch’s t test). Scale bar = 100µm. **(E)** Images of CNV lesions from aged WT and miR-34a^-/-^ mice. **(F)** Quantification of CNV lesions sizes for aged WT and miR-34a^-/-^ mice (n = 5 mice/group, unpaired Welch’s t test). Scale bars = 100µm.

### Disease relevance for miR-34a, KLF2, and CXCR4/CXCL12 in human choroidal vascular endothelial cells

To confirm the functional relevance of this pathway in neovascular lesions from patients with wet AMD, we stained CNV tissue samples from three different human donors via in situ hybridization (Fig. 7A). These tissue samples consisted entirely of neovascular membranes that were excised from patients with wet AMD during surgery that was performed as part of their usual care prior to the availability of anti-VEGFa or any other therapy including laser. We found miR-34a was abundantly expressed in human neovascular lesions from all three patients (Fig. 7B). We then used the interactive data viewer Spectacle (*72*) to comb through publicly available single-cell RNA sequencing data sets of human choroidal vascular endothelial cells in aging and with wet AMD. In one dataset (*73*), we found *KLF2* was highly expressed in a majority of cell types in the choroid, including vascular endothelial cells from veins, arteries, and choriocapillaris in young donors. In this same dataset, we found *KLF2* expression significantly decreased with age in all choroidal vascular endothelial cell types (Fig. 7C). This finding is especially important given that *KLF2* has been implicated in human aging and age-related vascular diseases (*57*, *74*). Next, we analyzed another single-cell dataset containing choroidal tissue of aged humans with or without wet AMD (*75*). We found *CXCL12* was upregulated in the choroidal arteries but downregulated in choroidal veins (Fig. 7D), potentially highlighting the role of arteries rather than veins as drivers of AMD development. Although *CXCL12* expression changed in opposite directions in the arteries and veins of patients with AMD, this difference contributes to the growing body of evidence that arteries and veins play distinct roles in AMD pathology (*76*). While a previous study found both *CXCL12* and *CXCR4* were increased in eyes with wet AMD (*77*), our analyses did not reveal any differences in *CXCR4* expression, possibly due to the limited amount of patient samples and datasets currently available. In conclusion, we provide evidence that gene expression of miR-34a, *KLF2*, and *CXCL12* are significantly altered in choroidal vascular endothelial cells from donor tissue with wet AMD or age.

**Fig. 7.**
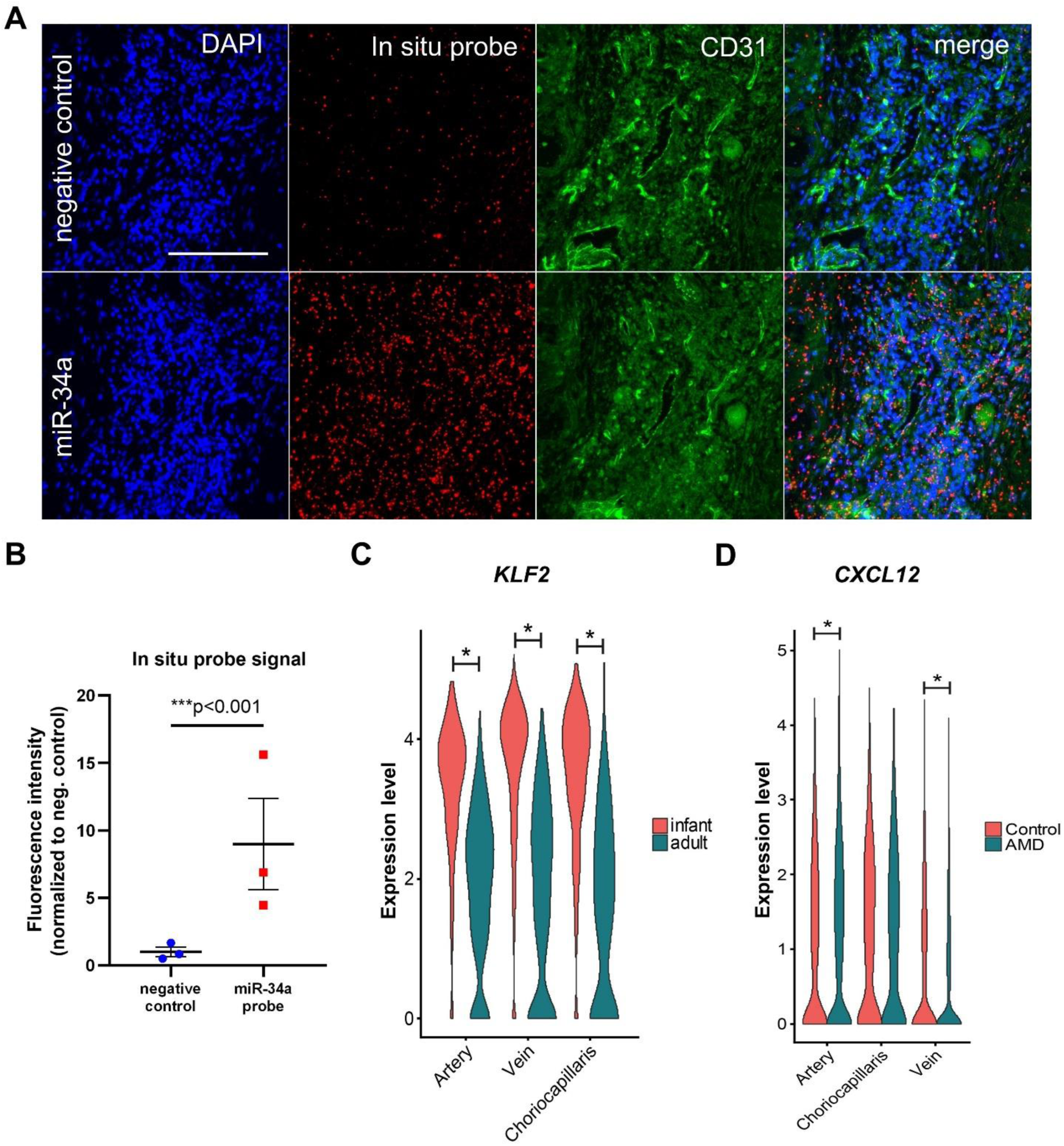
Human relevance for miR-34a, KLF2, and CXCL12 from human choroid tissue. **(A)** Sections of human CNV lesions dual-stained for miR-34a (red) via in situ hybridization and CD31 (green) via immunohistochemistry. Top row was stained using a scrambled control in situ hybridization probe, while the bottom row was stained with a human miR-34a-5p probe. Scale bar = 100µm. **(B)** Quantification of human CNV tissue samples from three different patients (n = 3 patient samples/group, ratio paired t test). **(C)** Violin plots showing KLF2 expression in each cell type (“*” denotes adjusted p-value<0.05). **(D**) Violin plots showing CXCL12 expression for each cell type (“*” denotes adjusted p-value<0.05 using the non-parametric Wilcoxon rank sum test).

Collectively, we show the downregulation of *KLF2* via miR-34a in vascular endothelial cells may promote pathological angiogenesis by facilitating the expression of *CXCR4* and *CXCL12*, without altering *VEGFa* or *KDR* expression. The data presented here corresponding to both human and mouse tissues provide translational relevance for the miR-34a/KLF2/CXCR4/CXCL12 pathway in pathological angiogenesis as seen in age-related ocular disease. (Fig. 8).

**Fig. 8.**
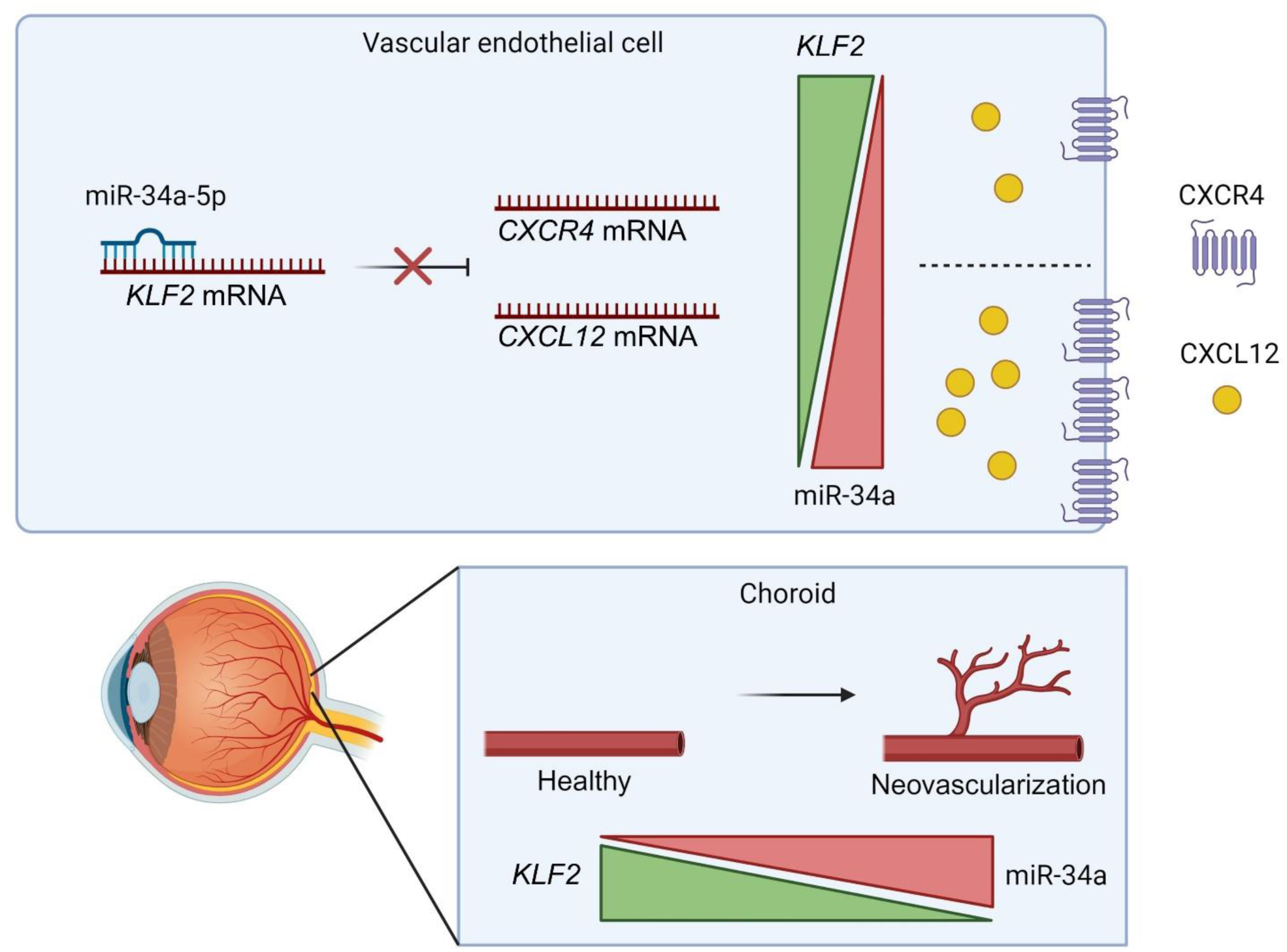
MiR-34a modulates pathological angiogenesis through the KLF2/CXCR4/CXCL12 pathway. MiR-34a directly inhibits KLF2 mRNA in vascular endothelial cells. The downregulation of KLF2 results in increased levels of CXCR4 and CXCL12 mRNA transcripts and proteins. The balance between miR-34a and KLF2 may regulate pathological angiogenesis as seen in patients with wet AMD. Created with BioRender.com (agreement number: RX277NTP8D).

## Discussion

Here, we investigated the role of miR-34a in the context of pathological angiogenesis in the choroid. First, we found that miR-34a was upregulated in vascular endothelial cells within CNV lesions of mice. Next, we determined that ablating miR-34a specifically in vascular endothelial cells was sufficient to reduce CNV lesion sizes. Then, by using HUVECs and staining murine CNV lesions we discovered that the presence or absence of miR-34a did not alter VEGFa or KDR expression in vascular endothelial cells. Using an integrative transcriptomic approach, we identified *KLF2* as a direct target of miR-34a in vascular endothelial cells. Our analyses also identified CXCR4 and CXCL12, pro-angiogenic factors regulated by KLF2 (*59*, *60*, *78*), to be upregulated in CNV lesions and HUVECs treated with miR-34a mimics. By inhibiting *KLF2*, we showed that *CXCR4* and *CXCL12* expression increased in human vascular endothelial cells, while not altering *VEGFa* or *KDR* expression. Additionally, suppression of *Klf2* in miR-34a^-/-^ mice promoted angiogenesis and reversed their small CNV phenotype. Afterwards, we repeated our CNV lesion size experiments and found that miR-34a also promotes CNV in aged mice. Next, we demonstrated that miR-34a was expressed in neovascular lesions excised from wet AMD patients. Finally, we analyzed single-cell data of human choroidal tissue and found that in vascular endothelial cells, *KLF2* expression decreased with age while *CXCL12* expression increased with wet AMD.

Our results highlight the proangiogenic features of miR-34a in ocular pathological angiogenesis. This finding is intriguing, especially in light of previous studies suggesting that miR-34a is anti-angiogenic in certain types of vascular endothelial cells and cancer cell lines, suggesting that downstream effects of miR-34a are tissue-dependent and influenced by microenvironmental cues (*79–83*). It is also crucial to highlight that these previous studies used drug interventions or conditioned media from cancer cells, which are likely to have additional impacts on gene expression. Conversely, another study found the upregulation of miR-34a in glioma cells promoted angiogenesis (*84*). The context-, tissue-, and cell-dependent roles for miR-34a in angiogenesis cannot be overstated, as it is predicted to inhibit over 700 different genes according to the miRbase database (*50*). Among these genes, *NOTCH1* (*85*), *VEGFa* (*86*), and *SIRT1* (*79*) can regulate angiogenesis. While both *NOTCH1* and *SIRT1* were significantly downregulated in our RNA-seq dataset of HUVECs treated with miR-34a mimics, these genes were not significantly altered in our analysis of the single-cell RNA-seq of lasered mouse choroids (Supplemental Data File 1), suggesting they may not play direct roles in CNV pathology. Our findings, in conjunction with the previous studies mentioned here, provide evidence that miR-34a can have pro-angiogenic properties depending on the cell type and context.

We demonstrate that miR-34a may promote pathological angiogenesis in a manner that is independent of VEGFa and its receptor KDR. Although *KDR* was significantly upregulated in our RNA-seq data set from HUVECs after miR-34a transfection, this finding was not validated via qPCR. Additionally, relative fluorescence intensity of Kdr staining was not altered in laser CNV lesions between miR-34a^-/-^ and WT mice. Furthermore, suppressing *KLF2 in vitro* did not influence *VEGFa* transcript expression in HUVECs or MVECs. Suppression of *Klf2* upregulated *Kdr* in MVECs, while it did not alter *KDR* transcript expression levels in HUVECs. While KLF2 has been shown to directly inhibit the *KDR* promoter (*51*), it remains to be seen if this relationship is relevant for our vascular disease model where both *Klf2* and *Kdr* were downregulated (Supplemental Data File 1), or in patients with wet AMD. This work provides evidence for an additional pathway to treat pathological angiogenesis, such as that in elderly patients with wet AMD, particularly by developing therapies that either upregulate *KLF2* and/or suppress the *CXCR4*/*CXCL12* axis.

We propose a role for miR-34a in modulating the KLF2/CXCL12/CXCR4 pathway in vascular endothelial cells, which in turn, contributes to pathogenic neovascularization. Notably, CXCL12 serves as a principal mobilization factor for endothelial progenitor cells (EPCs) (*87*), which actively participate in CNV (*88–90*). EPCs are derived from bone marrow and circulate in the bloodstream, where they contribute to the repair and regeneration of damaged vascular tissues and play a critical role in neovascular processes (*91*). Interestingly, miR-34a becomes upregulated in EPCs in response to increased shear stress from blood flow, promoting their differentiation into vascular endothelial cells (*92*). While our findings provide initial insights demonstrating how miR-34a, through its direct inhibition of KLF2, regulates CNV via the CXCR4/CXCL12 axis, the exact cellular contributions and mechanisms remain to be fully elucidated. It is plausible that the modulation of CXCL12 and CXCR4 by miR-34a-mediated suppression of KLF2 could influence not only local resident endothelial cells, but also the recruitment and function of EPCs, potentially enhancing the therapeutic efficacy of interventions targeting this pathway. Further detailed studies into the specific interactions between miR-34a, KLF2, CXCL12, and CXCR4 in both resident endothelial cells and circulating EPCs are necessary to fully understand their roles in the context of CNV and will be critical to develop targeted therapeutic strategies.

One area not explored in this investigation is whether the immune system is involved in miR-34a-related pathological angiogenesis. While we have previously demonstrated that immune cells play significant roles in regulating pathological angiogenesis in the eye (*17*, *33*, *34*), we did not observe miR-34a expression in immune cells in CNV lesions (Fig. S1, B to D). Of note, miR-34a promotes the pro-inflammatory M1 phenotype while suppressing M2 activation in macrophages (*93*), whereas *KLF2* suppresses inflammation and promotes the M2 phenotype in myeloid cells (*94–97*). Moreover, *KLF2* regulates an inflammatory response in HUVECs (*98*). We showed a similar inverse relationship between miR-34a and *KLF2* here in our study, where miR-34a upregulated the *CXCR4*/*CXCL12* axis while *KLF2* suppressed it. Additionally, the binding of CXCR4 to its canonical ligand CXCL12 directs the migration and adhesion of immune cells to inflammatory sites (*99*). This directional relationship between miR-34a and CXCR4/CXCL12 was also found in our study. For instance, *CXCR4* and *CXCL12* expression increased in miR-34a-treated HUVECs while their protein expression decreased in CNV lesions of miR-34^-/-^ mice. Future experiments could examine how miR-34a influences the recruitment of immune cells to CNV lesions and their adhesion to blood vessels.

Another area that warrants future investigation is whether any of the six other genes identified in our transcriptomic approach (Fig. 3A: *S1PR3*, *SGPP1*, *EDNRB*, *PITPNC1*, *RARB*, and *IVNS1ABP*) influence miR-34a-related pathological angiogenesis. Although *S1PR3* promotes angiogenesis through sphingosine-1-phosphate (*S1P*) (*100*), it was downregulated by miR-34a, so it is unlikely to be responsible for the decreased neovascularization seen in miR-34a^-/-^ mice. SGPP1 is a protein that recycles S1P, and although its expression is correlated with more angiogenesis, it is unclear whether decreased SGPP1 expression is pro-or anti-angiogenic (*101*). Additionally, *EDNRB* (*102*), *PITPNC* (*103*), and *RARB* (*104*) may all help to facilitate angiogenesis, so they are also unlikely to be negative regulators of CNV. Lastly, it is unclear if *IVNS1ABP* plays any role in angiogenesis.

In summary, we have shown that the upregulation of miR-34a in CNV lesions promotes pathological angiogenesis. We elucidated the molecular mechanism by which miR-34a regulates CNV within vascular endothelial cells in a VEGF-a independent manner. This study provides a better understanding of mechanisms of pathological angiogenesis and highlights potential areas of focus for future therapeutic interventions, namely KLF2, CXCR4, and CXCL12. Therapies designed against these factors, and used synergistically with current anti-VEGFa therapies, have the potential to reduce treatment burden and improve visual outcomes.

## Materials and Methods

### Mice

Mice were housed in approved animal facilities on a 12:12 light/dark schedule with ad libitum access to food and water, as well as daily veterinarian welfare checks and weekly cage changing. All mice were purchased from Jackson Labs, including miR-34a^-/-^ mice (*37*) (JAX stock #018279), miR-34a*^fl/fl^* mice (*38*) (JAX stock #018545), and VE-Cadherin-Cre mice (*39*) (JAX stock #006137). The Vglut2-ires-Cre-knock-in (*40*) (JAX stock #028863) was kindly donated by Dr. Phil Williams at Washington University in St. Louis. We crossed miR-34a*^fl/fl^* mice with VE-Cadherin-Cre and Vglut2-ires-Cre-knock-in mice to generate cell type-specific miR-34a-knock-out mice. All mice used in this study were female, 2-4 months of age for young mice, and 18-20 months for aged mice. All mice were maintained on a C57BL/6J background. When appropriate, mice in control and experimental groups were age matched. All animal experiments were approved by the IACUC of Washington University in St. Louis and were fully compliant with all applicable regulations and guidelines.

### Laser CNV

We conducted laser induced CNV following established procedures (*31*). Briefly, we anesthetized mice using a combination of ketamine and xylazine, and dilated pupils using topical tropicamide. Four CNV lesions were created around the optic disc (210 mW, 0.1 second, 100 μm spot size) using a slit-lamp delivery system with a cover glass as a contact lens. Following laser treatment, mice received topical antibiotic ointment applied to the ocular surface, were placed on a heating pad, and monitored until they regained consciousness from anesthesia. Mice were then housed in an IACUC approved facility where animal welfare checks occurred daily. Seven days following laser injury, mice were perfused with 100 μl of 50 mg/ml FITC-dextran (molecular weight, 2,000,000 daltons; MilliporeSigma) via the left femoral vein. Subsequently, eyes were enucleated and fixed in formalin for one hour at room temperature. After fixation, eyes were washed with PBS and flat-mounted the RPE/choroid complex onto a glass slide. We captured images of CNV lesions using a Leica DMi8 inverted microscope and excluded any CNV lesions with any signs of hemorrhaging. Finally, we quantified pixel intensity using MetaMorph (Molecular Devices).

### RNAscope In Situ Hybridization

To evaluate miR-34a expression, we used the RNAscope miRNAscope kit by ACD bio according to the manufacturer’s recommendations. Fresh-frozen whole-eye sections containing CNV lesions were fixed in 10% neutral buffered formalin at 4C for 1 hour. Slides were washed with increasing concentrations of ethanol at room temperature to dehydrate, then placed at -20C for no more than one week. Sections were selected with a hydrophobic marker and treated with hydrogen peroxide followed by protease IV for 15 minutes at room temperature in a humidified chamber. Slides were washed twice in PBS and then treated with the miR-34a probe and placed into a humidified oven at 40C for 2 hours. Following probe hybridization, the signal was boosted with a series of amplification steps. Finally, slides were stained with the fast red solution and DAPI. Slides were then mounted with the FluorSave Reagent.

### Immunostaining of choroid flat mounts

After sacrificing and enucleating eyes of lasered mice, eyes were fixed in 10% neutral buffered formalin at room temperature for 1-2 hours. Under a dissection microscope, the cornea, lens, retina, and optic nerves were removed leaving behind an intact choroid cup containing CNV lesions. Choroid cups were placed in blocking solution (PBS with 5% goat serum and 0.3% triton X) at 4C overnight. The next day, primary antibodies were added at the desired concentrations and choroid cups were placed back at 4C overnight. The following day, choroids were washed in blocking solution for 1 hour at room temperature for three times, then the secondary antibody and DAPI was added in blocking solution at 4C to incubate overnight. On the last day, choroid cups were washed three times in blocking solution for 1 hour each. Four cuts were made in the choroid cups in order to flatten the tissue and mount on slides using FluorSave reagent. The following primary antibodies were used: anti-VEGFa (abcam ab52917), anti-KDR (Invitrogen MA5-15157), anti-KLF2 (Bioss BS2772R), anti-CXCR4 (abcam ab18020), and anti-CXCL12 (abcam155090).

### Cell culture and microRNA mimic

We purchased primary HUVECs (PCS-100-010) from ATCC and cultured them in Vascular Cell Basal Medium (PCS-100-030) with Endothelial Cell Growth Kit (PCS-100-041). Cells were maintained subconfluently and used before passage 4 for all experiments. MiR-34a and negative control mimics (Qiagen miRCURY LNA miRNA Mimics) were transfected into HUVECs using Lipofectamine RNAiMAX (Invitrogen 13778100) according to the manufacturer’s recommendations.

### Knock-down of *Klf2*

Premium LNA Gapmers targeting *Klf2* were designed and purchased from Qiagen and transfected into HUVECs and MVECs with Lipofectamine 3000 (Invitrogen L3000001) according to the manufacturer’s recommendations. For *in vivo* experiments, mice received daily intraperitoneal injections of 10mg/kg *in vivo* LNA *Klf2* or negative control in sterile PBS starting on the day of laser injury and ending seven days later.

### RNA and miRNA isolation

We isolated RNA from HUVECs or MVECs 48 hours after transfection cells were trypsinized, neutralized, and spun down in an Eppendorf tube at 250 g for 3 min to remove any dead cells resulting from the transfection. Cells were resuspended in Buffer RLT lysis buffer with 1% Beta-Mercaptoethanol. We then followed the manufacturer’s instructions for RNA isolation using the RNeasy Mini Kit (Qiagen 74104). RNA yield was measured on a Tecan NanoQuant Plate.

We isolated miRNA using the mirVana miRNA Isolation Kit, with phenol (ThermoFisher Scientific AM1560) according to the manufacturer’s instructions.

### Bulk RNA sequencing analysis

After RNA isolation, samples were sent to the Genome Technology Access Center (GTAC) at Washington University School of Medicine in St. Louis for further processing. Total RNA integrity was determined using Agilent Bioanalyzer or 4200 Tapestation. Library preparation was performed with 500 ng to 1 ug of total RNA. Ribosomal RNA was removed by an RNase-H method using RiboErase kits (Kapa Biosystems). mRNA was then fragmented in reverse transcriptase buffer and heating to 94 degrees for 8 minutes. mRNA was reverse transcribed to yield cDNA using SuperScript III RT enzyme (Life Technologies, per manufacturer’s instructions) and random hexamers. A second strand reaction was performed to yield ds-cDNA. cDNA was blunt ended, had an A base added to the 3’ ends, and then had Illumina sequencing adapters ligated to the ends. Ligated fragments were then amplified for 12-15 cycles using primers incorporating unique dual index tags. Fragments were sequenced on an Illumina NovaSeq-6000 using paired end reads extending 150 bases. Basecalls and demultiplexing were performed with Illumina’s bcl2fastq software with a maximum of one mismatch in the indexing read. RNA-seq reads were then aligned to the Ensembl release 101 primary assembly with STAR version 2.7.9a. Gene counts were derived from the number of uniquely aligned unambiguous reads by Subread:featureCount version 2.0.3. Isoform expression of known Ensembl transcripts were quantified with Salmon version 1.5.2. Sequencing performance was assessed for the total number of aligned reads, total number of uniquely aligned reads, and features detected. The ribosomal fraction, known junction saturation, and read distribution over known gene models were quantified with RSeQC version 4.0.

### Neovascular AMD patient samples

Lesions were excised from three different patients with neovascular AMD as part of their usual care, prior to anti-VEGFa or photocoagulation treatment. These samples were excised during sub-macular surgery and preserved in a de-identified manner in paraffin blocks. The samples consist exclusively of neovascular endothelial tufts; they are not eye cross-sections but are instead specific tissues removed during surgery. The Human Research Protection Office at Washington University in St. Louis has deemed these samples exempt from informed consent, under HRPO#09-1058/Apte.

### Analysis of single cell RNA sequencing data

We downloaded three previously published single cell RNA sequencing datasets: 1) of murine CNV lesions (*49*); 2) of human infant vs. adult choroid (*73*); and 3) of human choroid in control vs. AMD patients (*75*). For each of these datasets, we downloaded the raw count matrices and re-processed the data using Seurat v4 (*105*) to normalize, log-transform, and scale the data; and then identified the top 2,000 highly variable genes for principal component analysis. We next used the Harmony package to integrate the data across each sample (*106*). We performed differential gene expression analysis for each of the three datasets as follows.

#### Murine CNV lesions

To identify marker genes of CNV-induced endothelial cell states, we performed differential gene expression analysis using Seurat’s built-in FindMarkers() function. We compared 5 pathologic endothelial cell states including proliferating endothelial cells, tip cells, CNV transitioning cells, immature endothelial cells, and neophalanx (n=1,966 cells) to all other non-pathologic endothelial cell states in the dataset (n=26,371 cells). We used Seurat’s FindMarkers() function to identify any differentially expressed genes with |log2(fold-change)| > 0.25 and adjusted p-values < 0.05.

#### Infant vs. adult choroid

We identified genes that were differentially expressed in infant (samples 24, 25) vs. aged choroids (19, 21, 22, 23). We performed this analysis in a cell population-specific manner with sample sizes ranging from n=119 to 7,286 cells. We identified any differentially expressed genes with adjusted p-values < 0.05.

#### Control vs. AMD choroid

We identified genes that were differentially expressed in control (samples 1, 2, 3, 4, 5, 7, 8, 9, 6, 10) vs. AMD (samples 11, 12, 13, 14, 15, 16, 17, 18, 19, 20, 21). We performed this analysis in a cell population-specific manner with sample size ranging from n=109 to 4,519. We identified any differentially expressed genes with adjusted p-values < 0.05.

### cDNA synthesis and qPCR

Total RNA collected from HUVECs and MVECs were reverse transcribed to cDNA using the high-capacity reverse transcription kit (Applied Biosystems 4368814) according to the manufacturer’s instructions. We then used the TaqMan fast advanced master mix (Thermo Fish Scientific 4444557) to perform qPCR. We normalized gene expression to actin, and we used the ΔΔCT method to analyze the data. The following TaqMan probes were used for experiments involving human cells: *Actin* (Hs01060665_g1), *VEGFa* (Hs00900055_m1), *KDR* (Hs00911700_m1), *KLF2* (Hs00360439_g1), *CXCR4* (Hs00607978_s1), *CXCL12* (Hs03676656_mH). The following TaqMan probes were used for experiments involving mouse cells: *Actin* (Mm02619580_g1), *Vegfa* (Mm00437306_m1), *Kdr* (Mm01222421_m1), *Klf2* (Mm00500486_g1) *Cxcl12* (Mm00445553_m1), *Cxcr4* (Mm01996749_s1).

For measuring miR-34a transcript levels, we used the miRCURY LNA miRNA PCR Assay (Qiagen). miR-34a gene expression was normalized to U6 and data was analyzed using the ΔΔCT method.

### Dual luciferase assay

We ordered custom *KLF2* 3′ UTR miRNA target clones from GeneCopoeia with wild-type or mutated *KLF2* 3′ UTR sequences inserted downstream of a secreted GLuc reporter gene driven by the SV40 promoter and a secreted alkaline phosphatase (SEAP) reporter gene driven by a CMV promoter. Reporter plasmids were dual-transfected into HUVECs with miR-34a mimics or negative control mimics using Lipofectamine 3000 (Invitrogen L3000001) according to the manufacturer’s recommendations. Supernatant was collected 48 hours after transfection and Gluc activity was measured with the Secrete Pair Dual Luminescence assay kit (GeneCopoeia LF031). Transfection efficiency was normalized by measuring SEAP activity.

## Funding

National Institutes of Health R01 EY019287 (RSA)

Jeffrey T. Fort Innovation Fund (RSA)

Starr Foundation AMD Research Fund (RSA)

Siteman Retina Research Fund (RSA)

Carl Marshall and Mildred Almen Reeves Foundation (RSA)

Retina Associates of St. Louis Research Fund (RSA)

Research to Prevent Blindness/American Macular Degeneration Foundation Catalyst

Award for Innovative Research Approaches for Age-Related Macular Degeneration (RSA)

National Institutes of Health P30 EY02687 (Vision Core Grant)

An unrestricted grant from Research to Prevent Blindness to the John F. Hardesty, MD

Department of Ophthalmology and Visual Sciences at Washington University School of Medicine in St. Louis.

Bayer Retina Award in Japan (RT)

The International Retinal Research Foundation (RT)

The Japan Society for the Promotion of Science Overseas Research Fellowship (RT)

National Institutes of Health Training Grant (1T32GM1397740-1) (TJL)

National Institutes of Health Training Grant (F30DK130282) (JBL)

Washington University in St. Louis Medical Scientist Training Program (National Institutes of Health Training Grant T32GM07200) (JBL)

We acknowledge the Genome Technology Access Center at the McDonnell Genome Institute at Washington University School of Medicine for help with genomic analysis.

The Center is partially supported by NCI Cancer Center Support Grant #P30 CA91842 to the Siteman Cancer Center from the National Center for Research Resources (NCRR), a component of the NIH, and NIH Roadmap for Medical Research.

## Author contributions

Conceptualization: JJC, RSA

Methodology: JJC, RSA, JBL

Visualization: JJC, JBL

Investigation: JJC, JBL, RT, TJL, AS

Writing—original draft: JJC

Writing—review & editing: JJC, RSA, JBL, RT, TJL, AS

## Competing interests

Authors declare that they have no competing interests.

## Data and materials availability

Raw sequencing files for bulk RNA-seq experiments are deposited in GEO with the following accession number: GSE252957. All data are available in the main text or the supplementary materials.

## Supplementary Materials

**Fig. S1.**
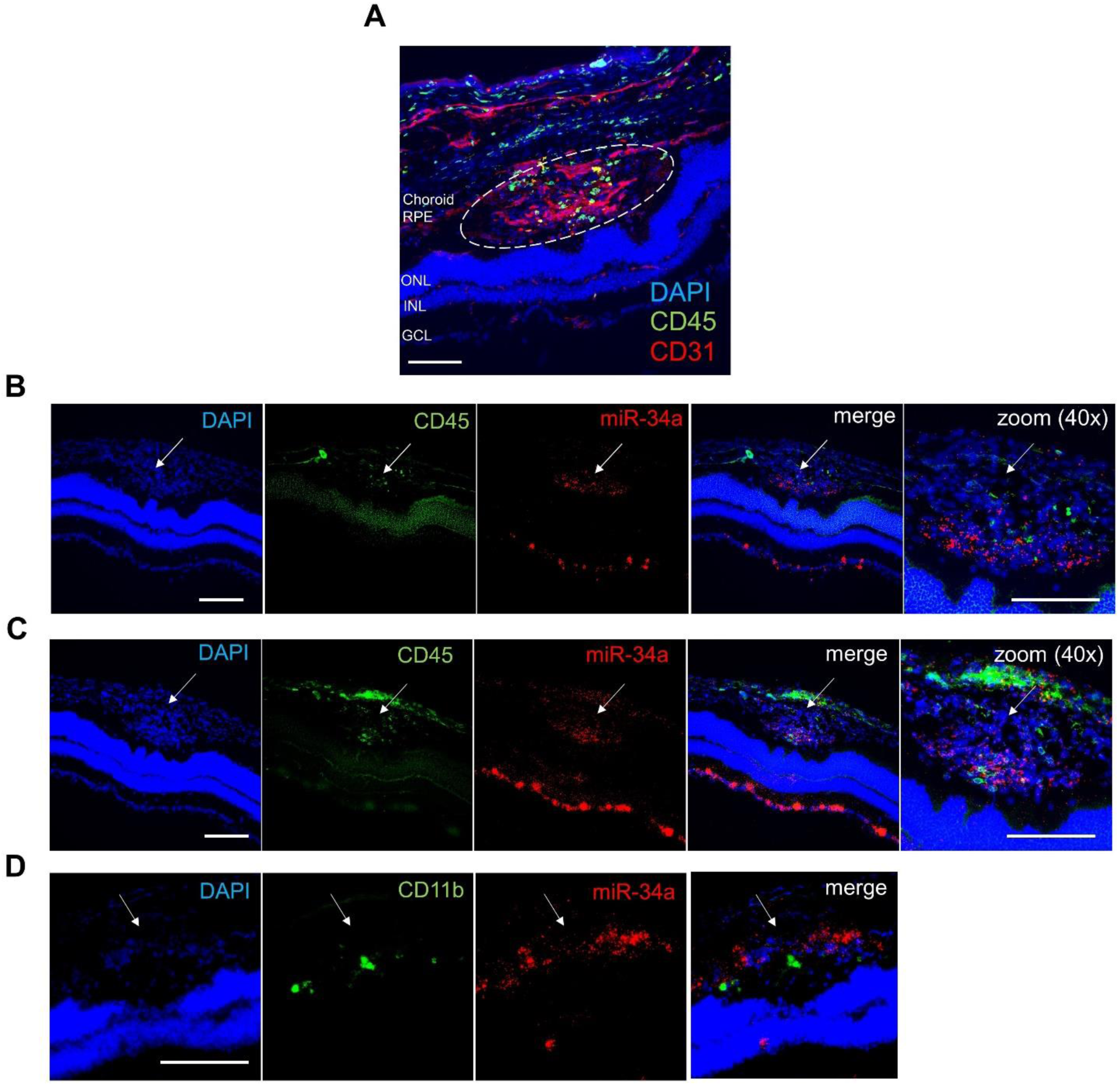
MiR-34a expression does not overlap with immune cells in murine CNV. **(A)** Image of a CNV lesion from a WT mouse stained for the general immune cell marker CD45 and the vascular endothelial cell marker CD31. **(B)** Image of a CNV lesion from a WT mouse co-stained with CD45 and miR-34a. **(C)** Image of a CNV lesion from a WT mouse co-stained with CD45 and miR-34a. **(D)** Image of a CNV lesion from a WT mouse co-stained with the myeloid-lineage marker CD11b and miR-34a. RPE denotes retinal pigment epithelium, ONL denotes outer nuclear layer, INL denotes inner nuclear layer, and GCL denotes ganglion cell layer. Scale bars = 100µm.

**Fig. S2.**
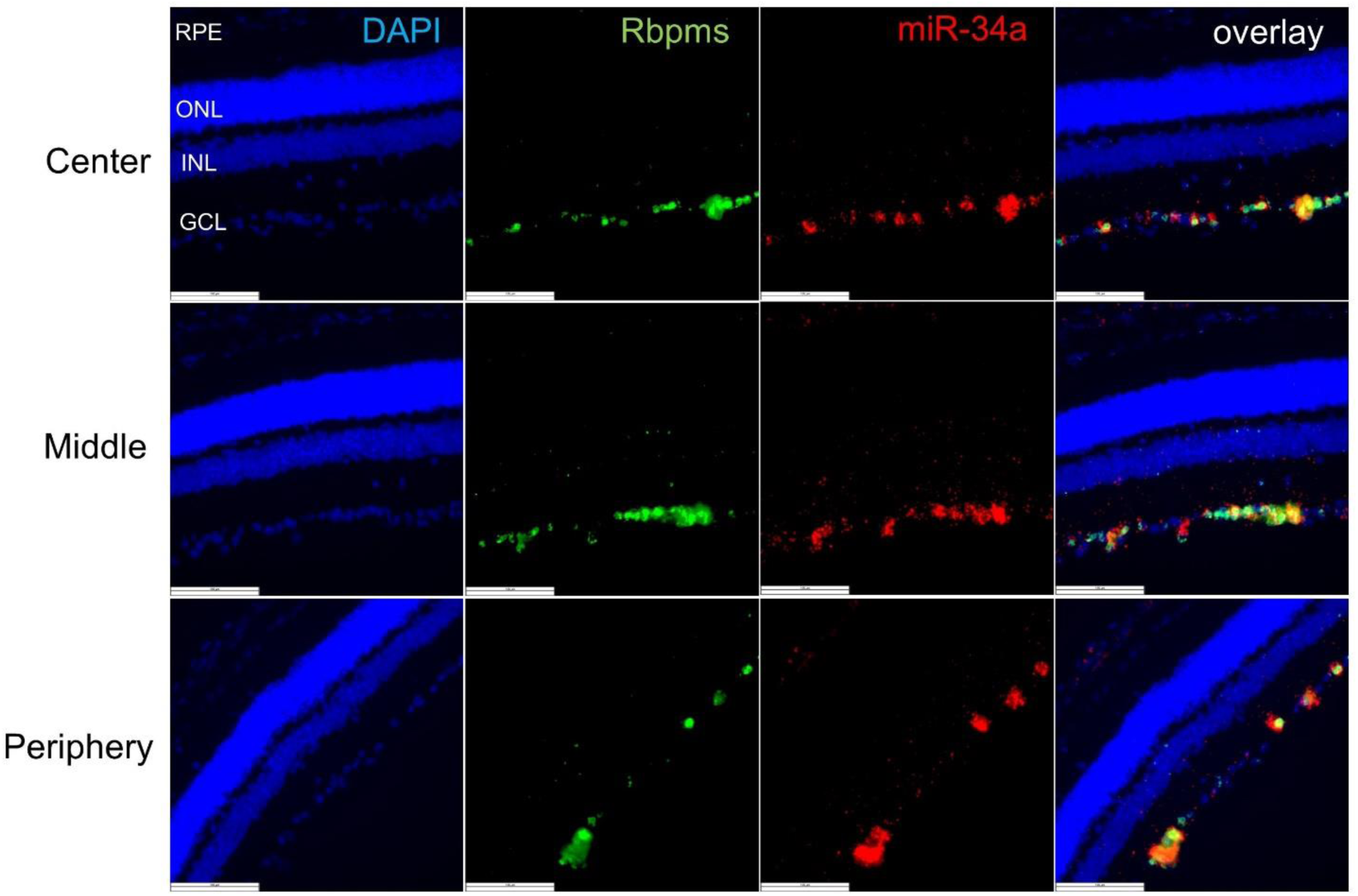
MiR-34a is expressed in RGCs under normal conditions. Fresh-frozen whole eye sections from a WT mouse co-stained for the RGC marker RBPMS and miR-34a. RPE denotes retinal pigment epithelium, ONL denotes outer nuclear layer, INL denotes inner nuclear layer, and GCL denotes ganglion cell layer. Scale bar = 100µm.

**Fig. S3.**
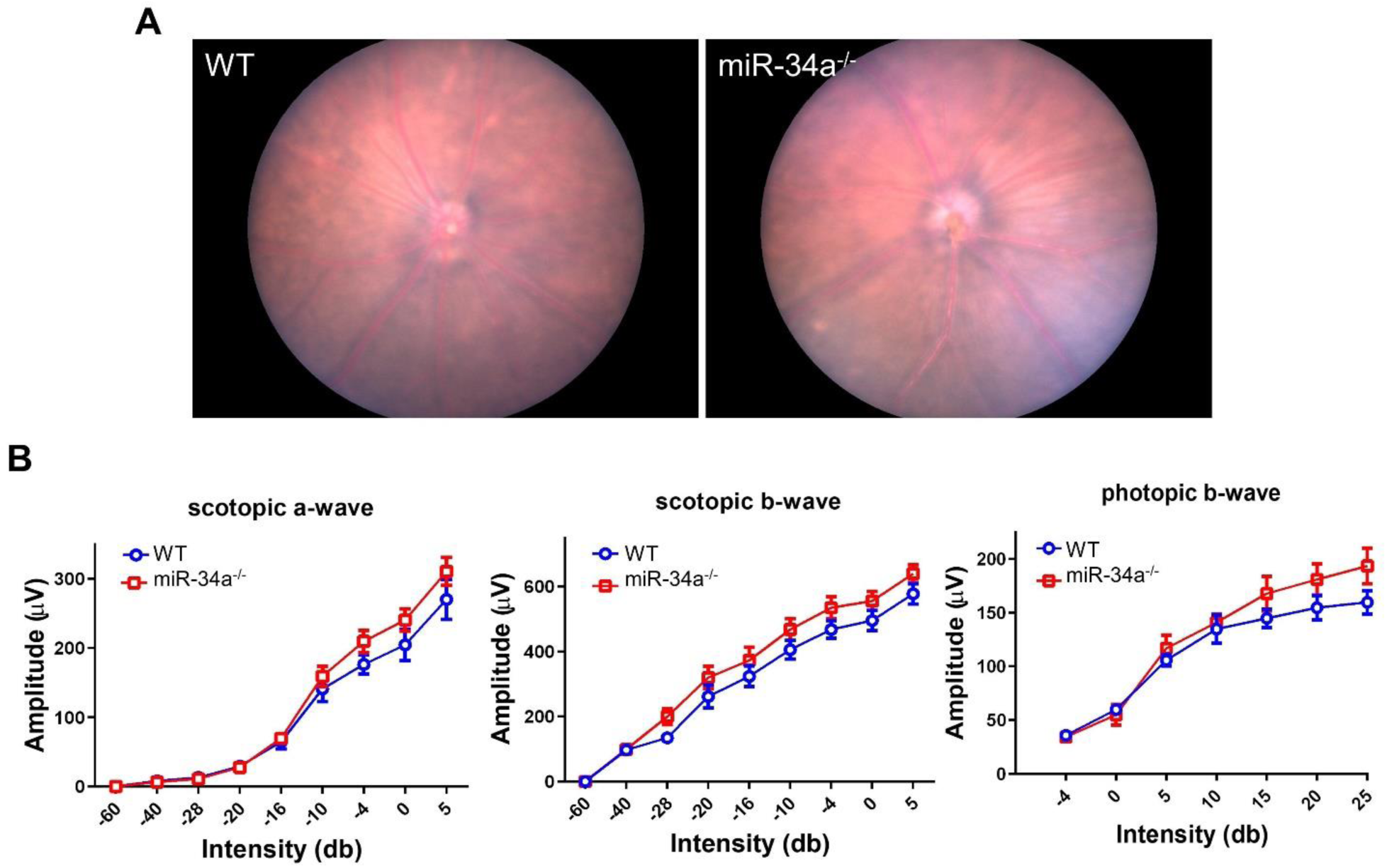
MiR-34a^-/-^ mice have no structural or functional ocular abnormalities as seen via fundoscopy and ERG. **(A)** Fundoscopy images of WT and miR-34a^-/-^ mice. **(B)** ERG data from WT and miR-34a^-/-^ mice (n = 4 mice/group, Two-way ANOVA with Bonferroni’s multiple comparison test).

**Fig. S4.**
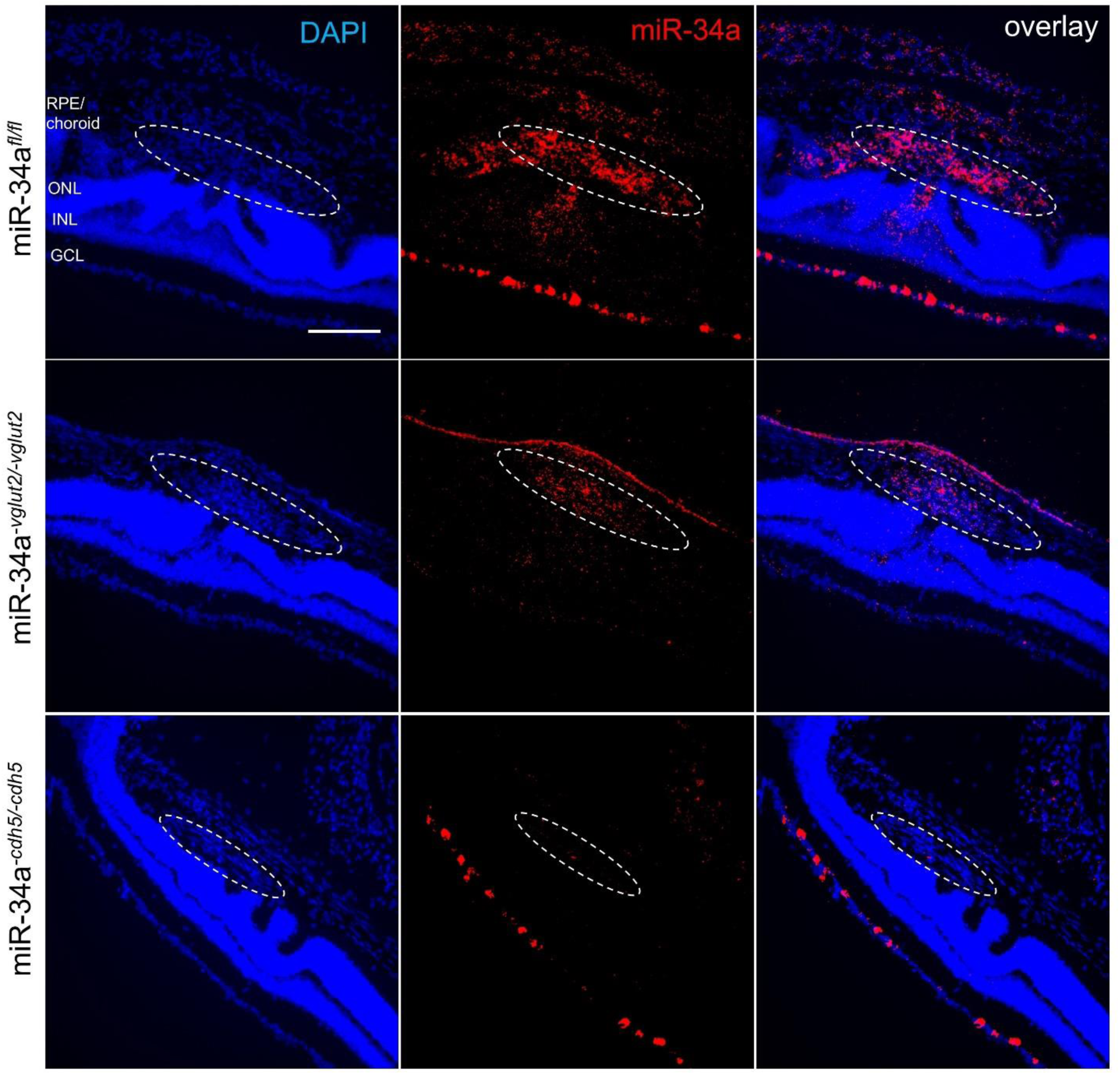
Validation of cell-type specific miR-34a knock-out mice. Top row: Cross-sectional images of injured miR-34a^fl/fl^ mouse eyes stained for miR-34a and DAPI, showing positive signal in both CNV lesions and RGCs. Middle row: Cross-sectional images of injured miR-34a^-vglut2/-vglut2^ mouse eyes stained for miR-34a and DAPI, showing positive signal in CNV lesions, but not RGCs. Bottom row: Cross-sectional images of injured miR-34a^-cdh5/-cdh5^ mouse eyes stained for miR-34a and DAPI, showing no signal in CNV lesions, but with positive signal in RGCs. RPE denotes retinal pigment epithelium, ONL denotes outer nuclear layer, INL denotes inner nuclear layer, and GCL denotes ganglion cell layer. Scale bar = 100µm.

**Fig. S5.**
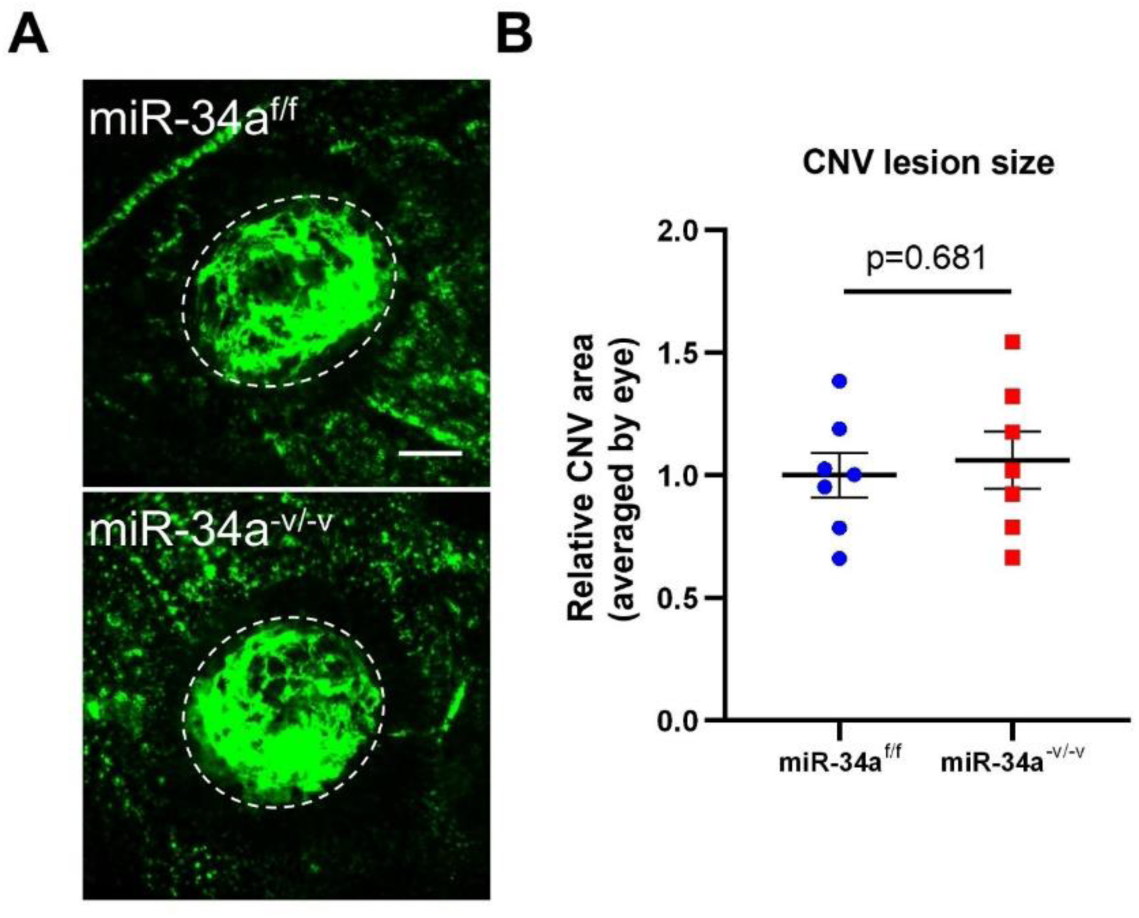
RGC-specific miR-34a expression does not influence CNV after laser injury. **(A)** Images of CNV lesion sizes between miR-34a^fl/fl^ (miR-34a^f/f^) and miR-34a^-vglut2/-vglut2^ (miR-34a^-v/-v^) mice. **(B)** Quantification of CNV lesions sizes between miR-34a^fl/fl^ and miR-34a^-vglut2/-vglut2^ mice (n = 4 mice/group, unpaired Welch’s t test). Scale bar = 100µm.

**Fig. S6.**
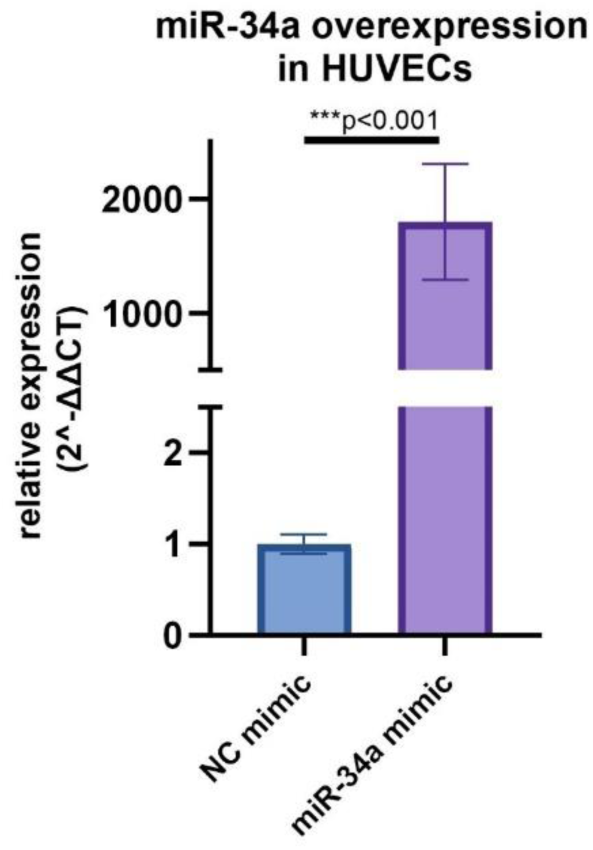
Successful overexpression of miR-34a in HUVECs. HUVECs were successfully transfected with either miR-34a or scrambled negative control mimics as confirmed via qPCR for miR-34a expression. (U6 was used an endogenous control, n = 3/group, unpaired Welch’s t test).

**Fig. S7.**
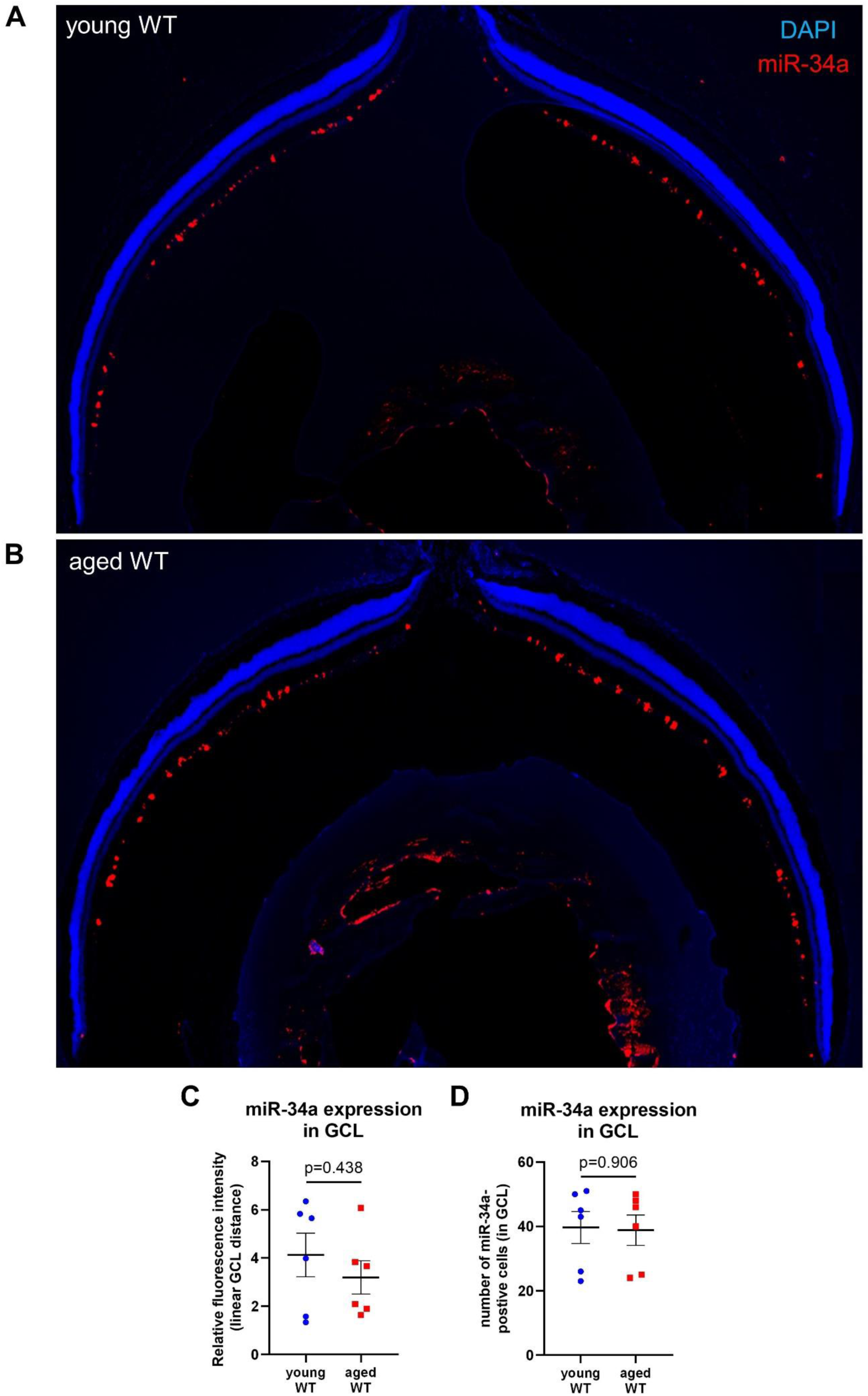
MiR-34a expression does not change with age in the mouse eye under homeostatic conditions. **(A, B)** Fresh-frozen whole-eye sections of young or aged WT mice stained for miR-34a. **(C)** Relative fluorescence intensity of miR-34a signal in the ganglion cell layer (GCL) (n = 6 mice/group, unpaired Welch’s t test). **(D)** Number of miR-34a positive cells in the GCL (n = 6 mice/group, unpaired Welch’s t test).

## Data S1. (separate file)

This Excel file contains the results of differential gene expression analysis from: 1) bulk RNA-sequencing of HUVECs transfected with miR-34a mimics compared to scrambled negative controls; and 2) single-cell RNA-sequencing of cells collected from murine laser-CNV samples comparing diseased vascular endothelial cells with healthy vascular endothelial cells.

## Notes

### Competing Interest Statement

The authors have declared no competing interest.

